# General predictions for the effects of warming on competition

**DOI:** 10.1101/2025.10.02.680125

**Authors:** Kaleigh E. Davis, Tess N. Grainger, Po-Ju Ke, Patrick L. Thompson, Mary I. O’Connor, Joey R. Bernhardt

## Abstract

Understanding the effects of climate change on ecological communities has been limited by a lack of general theory for how temperature affects competition. To fill this knowledge gap, we integrated Modern Coexistence Theory and the Metabolic Theory of Ecology by incorporating empirically-derived temperature sensitivities into Modern Coexistence Theory’s central model. We then simulated warming in consumer-resource systems and found that warming reduced both niche and fitness differences, making species more ecologically similar and competitive interactions more neutral. The greatest shifts in competition occurred when temperature sensitivities among species were highly asymmetrical. Effects of warming on competition via niche differences were comparable to those on fitness differences, suggesting that the emphasis on vital rates in global change research may overlook key biodiversity drivers. This general theory expands the domains of two prominent ecological theories and provides predictions for how warming may alter competition even in benign regions of species’ thermal niches.

## Introduction

Understanding the effects of climate warming on biodiversity is a central challenge of modern ecology (Parmesan & Yohe 2003; Peters & Lovejoy 1992; Román-Palacios & Wiens 2020). Species interactions, and competition in particular, influence species’ responses to the environment, and are thus critical to understanding the effects of warming on biodiversity (Godoy 2019; Tilman 1994). While a considerable body of empirical work has demonstrated shifts in competition with changing environments (e.g., Alexander *et al*. 2015; Germain *et al*. 2018; Pérez-Ramos *et al*. 2019), the nature of these shifts has been variable, and explanations usually context specific. Whether there are any general patterns in how warming affects competition, or whether these changes are inherently species-specific is a major open question. In the face of rapid environmental change, new theory is needed to address this question.

Modern Coexistence Theory has become the leading framework for understanding competition and coexistence (Chesson 2000; Godoy & Levine 2014). This theory has the advantage of being agnostic to the specific mechanisms or context of competitive interactions, which allows it to be used to investigate general principles of competition and to be applied across a wide range of systems (Godwin *et al*. 2020; Grainger *et al*. 2019). Within Modern Coexistence Theory, in the absence of environmental fluctuations, species’ competitive differences are described along two axes: fitness differences and niche differences (Chesson 2000). Fitness differences are interspecific differences that give one species a competitive advantage and stem from differences between competing species in intrinsic growth rate and sensitivity to competition. Niche differences are differences that reduce the strength of interspecific competition relative to intraspecific competition, for example, differences in resource use or susceptibility to predators (Chesson 2000; Chesson & Kuang 2008). The balance of these differences determines whether two species can coexist, one species will exclude the other, or they are competitively neutral (Adler *et al*. 2007). In global change research, work to understand the effects of warming on species abundance and distribution has predominantly emphasized shifts in vital rates (Spaak *et al*. 2021; fitness differences; e.g. Merow et al 2014, Ehrlen and Morris 2015; but see Adler et al., 2012; Usinowicz & Levine 2018), while the potential effects of warming on niche differences remain underexplored.

One criticism of Modern Coexistence Theory is that, because it is phenomenological, it is not well-suited to generate a priori predictions about how a given factor will affect competition and coexistence (Terry 2025). Indeed, the parameters used to estimate niche and fitness differences in classical Modern Coexistence Theory models describe the outcomes of competitive interactions but are not related explicitly to underlying mechanisms (Chesson 2000). However, niche and fitness differences can be alternatively calculated from their constituent mechanistic processes (Chesson & Kuang 2008; Letten *et al*. 2017), for example, parameters related to resource competition (Song *et al*. 2019). Fortunately, the effects of temperature on many of these mechanistic processes have also been investigated empirically (Dell *et al*. 2011; Rall *et al*. 2010). To evaluate how these temperature effects shift competitive dynamics with warming, the logical next step is to integrate empirically-grounded temperature dependencies into Modern Coexistence Theory models.

The Metabolic Theory of Ecology is a widely used framework for understanding the effects of temperature on ecological processes. A central premise of this framework is that ecological processes, such as population growth and species interactions, all stem from metabolic processes, which control the uptake and transformation of resources into maintenance, growth and reproduction (Brown *et al*. 2004). In this way, metabolism fundamentally and predictably constrains higher order processes across systems, leading to a predictable “scaling” of temperature effects across levels of organization. From this principle of metabolic scaling, the Metabolic Theory of Ecology has been used to predict the effect of temperature on population growth rates (Savage *et al*. 2004), population carrying capacities (Bernhardt *et al*. 2018a), consumer-resource interactions (O’Connor 2009; O’Connor *et al*. 2011), food webs (Gibert *et al*. 2022; Gilbert *et al*. 2014; Vasseur & McCann 2005), and ecosystem productivity (Enquist *et al*. 2003; Lopez-Urrutia *et al*. 2006; Welter *et al*. 2015). Despite this broad success, extending the metabolic scaling approach to competitive interactions has proven challenging, due to a conflict in focus between competition theory and the Metabolic Theory of Ecology.

While competition theory has focused primarily on changes in ecological outcomes driven by interspecific variation in common underlying traits (here, intra-process asymmetry, Fig. 1) (e.g., Kraft *et al*. 2008; Weiher *et al*. 1998), the Metabolic Theory of Ecology has largely ignored interspecific variation in thermal traits, and focused instead on average differences in thermal responses across metabolic processes (here, inter-process asymmetry, Fig. 1, but see Clegg & Pawar 2024; Gibert *et al*. 2022). Since our focal traits are thermal traits, we will refer to these intra- and inter-process thermal asymmetries (Fig. 1). Progress to incorporate interspecific variability in thermal traits into metabolic scaling studies of competition has been made primarily by exploring consequences of high, above optimal, temperature, and generalist-specialist trade-offs in thermal tolerance breadth (e.g., Amarasekare & Johnson 2017; Simon & Amarasekare 2024; Smith & Amarasekare 2018). However, most species live, on average, well below their thermal optimum in nature (Bernhardt *et al*. 2018b; Martin & Huey 2008), and whether interspecific trait differences in the non-stressful (sub-optimal) temperature range, alone, can produce general patterns in competitive dynamics under warming is an open question. Incorporating temperature-dependence at sub-optimal temperatures into the governing processes of Modern Coexistence Theory enables us to answer this question.

**Figure 1.**
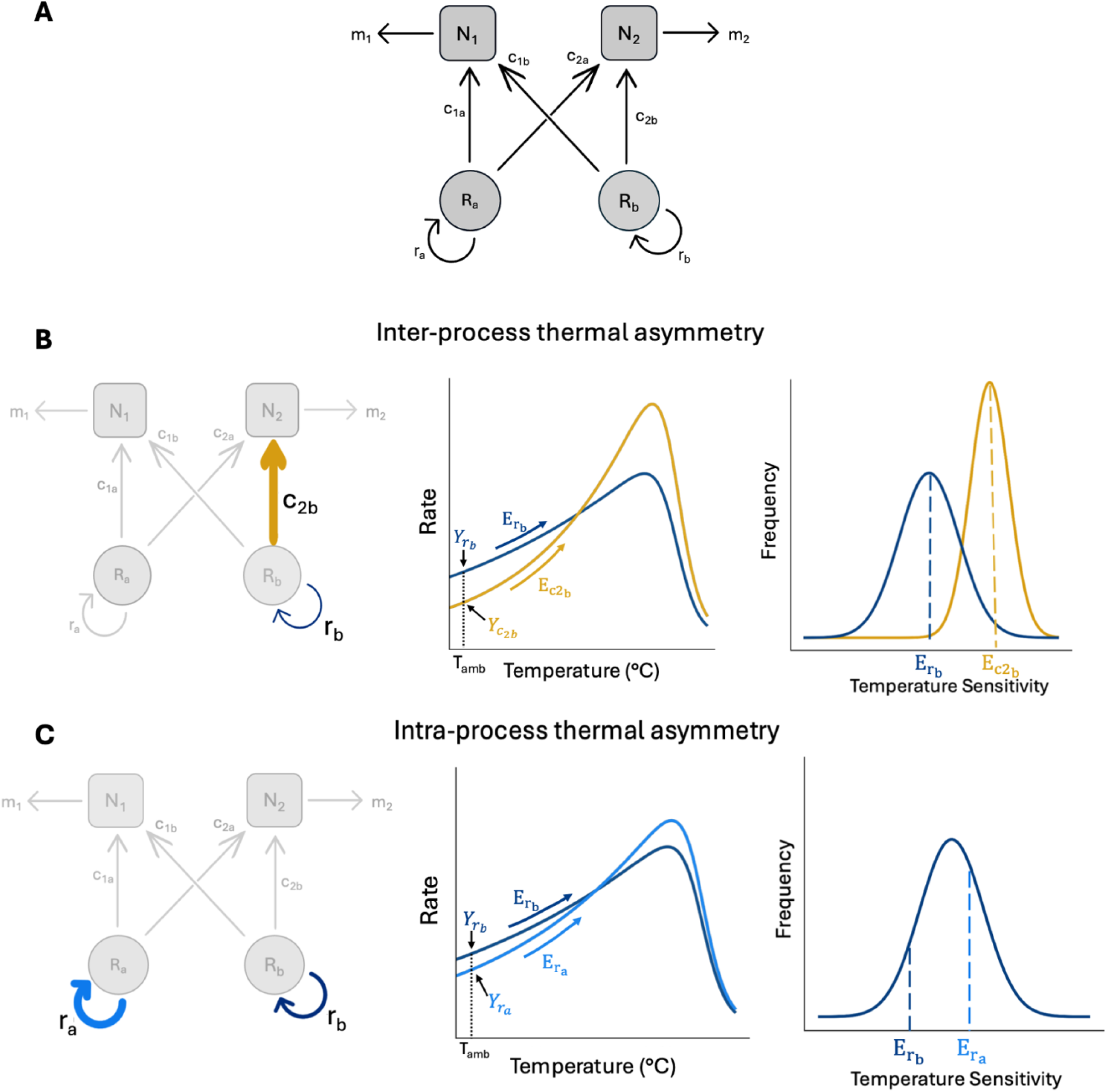
Schematic of a consumer-resource model with temperature dependent processes, yielding inter- and intra-process thermal asymmetries. In panel A, two unique consumers are shown as boxes (*N_1_* and *N_2_*) and two substitutable resources as circles (*R_a_* and *R_b_*). Arrows show directional energy flow processes and are labeled with the model parameters they represent: *m_i_* is the mortality of consumer *i*, *c_ik_* is consumption rate of resource *k* by consumer *i*, and *r_k_* is the intrinsic growth rate of resource *k*. Parameters *K_k_*, resource carrying capacity, and *v_ik_*, conversion efficiency, are not shown, but mediate the abundance of available resource, *R_k_*, and the transfer of energy from *R_k_* to *N_i_*, respectively. Panels B and C show how temperature sensitivity, *E*, or the increase in the rate of a process with temperature below the temperature where the curve peaks (central panels), is incorporated into the model via biological rates of the consumer and resource, and how this can generate two types of thermal asymmetries depending on where in the consumer-resource system a difference in temperature sensitivity occurs (left panels). The ambient temperature is given as *T_amb_*, and the rates of a process at this temperature, corresponding to 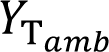 in Eq. 9, are given. The thickness of the arrows in the left panels corresponds to the temperature sensitivity depicted in the central panels. The right panels in B and C relate a single observation of temperature sensitivity (dashed line) for a parameter to the broader empirical distribution of temperature sensitivities for that parameter constructed from our data synthesis. In an inter-process thermal asymmetry (Panel B), the two temperature sensitivities come from independent distributions of temperature sensitivity in each parameter, whereas in an intra-process thermal asymmetry (Panel C), the two temperature sensitivities come from the same distribution of temperature sensitivity.

Here we draw from the Metabolic Theory of Ecology and Modern Coexistence Theory to develop theory for how warming affects competition for shared resources at suboptimal temperatures. We use this theory to address three questions: 1) How do the temperature sensitivities of each process underlying competition contribute to changes in niche and fitness differences with warming? 2) How do inter- and intra-process thermal asymmetries, affect these changes? 3) Are there general effects of warming on competition that emerge from the combined effects of warming on its suite of underlying mechanistic processes? To answer these questions, we analyzed a MacArthur consumer–resource model (MacArthur 1970) using the canonical two-species Lotka–Volterra formulation of Modern Coexistence Theory that expresses phenomenological competition coefficients with mechanistic parameters (Chesson 1990, 2011; Chesson & Kuang 2008; Song *et al*. 2019). We modeled the effects of warming on competition by making each parameter in the MacArthur consumer–resource model temperature-sensitive. We synthesized temperature sensitivity data for each model parameter from the published literature to generate an empirical distribution of temperature sensitivities and then used this distribution to simulate effects of warming on competition between species pairs. In doing so, we identified a novel set of general predictions for how competition is affected by warming, and the conditions under which we expect to see more extreme shifts in competition.

## Materials and Methods

### Resource competition model

We modeled competition in a two-consumer system (*N_1_* and *N_2_*) with two substitutable resources (*R_a_* and *R_b_*) following the classic MacArthur consumer–resource model (MacArthur 1970):

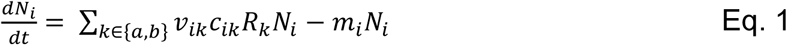

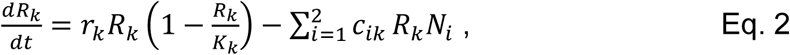

with *i* = *1* or *2* and *k* = *a* or *b* (see Fig. S1 for simulations with >2 resources). Here, the resource follows logistic growth with intrinsic growth rate 𝑟_𝑘_ and carrying capacity 𝐾_𝑘_. Consumer species *i* uses resource *k* following a Holling type-I functional response, with per capita consumption rate (hereafter “consumption rate”) parameter 𝑐_𝑖𝑘_ and conversion efficiency parameter 𝑣_𝑖𝑘_. While the type-I functional response is sometimes considered unrealistic, recent work suggests that it may be appropriate in more cases than previously thought (Coblentz *et al*. 2023; Novak *et al*. 2025). Finally, consumers experience per-capita mortality 𝑚_𝑖_. We converted this into a two-species Lotka–Volterra competition model by separation of timescales. Specifically, by assuming fast resource dynamics, the quasi-equilibrium of 𝑅_𝑘_ can be solved as:

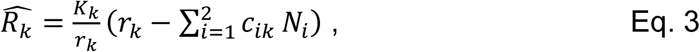

which is a function of consumer abundances. Substituting this expression back into Eq. 1, we arrive at the following:

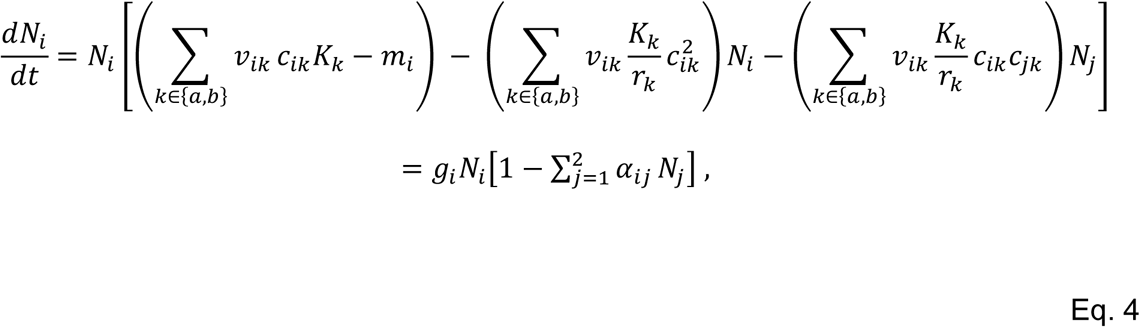

with

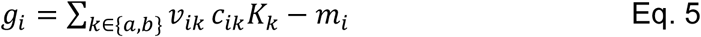

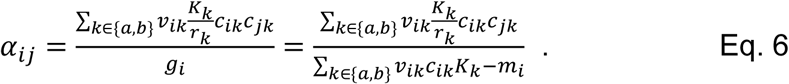

Here, 𝑔_𝑖_ represents consumer *i* ’s intrinsic growth rate, which equals the maximum energy gain by consumption minus the consumer’s per-capita mortality rate. On the other hand, 𝛼_𝑖𝑗_ represents the relative reduction in consumer species *i* ’s intrinsic growth rate caused by competition from species *j* (i.e., relative competition coefficient; sensu Song *et al*. 2019).

Modern Coexistence Theory then defines niche overlap (𝜌) and fitness ratio 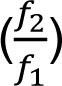 as follows:

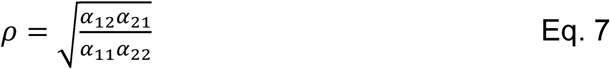

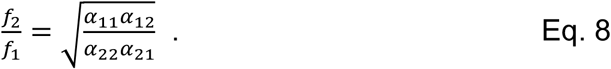

Here, 𝜌 quantifies the relative strength of intraspecific and interspecific competition, and 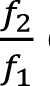 quantifies species’ relative sensitivity to competition. Finally, we define niche differences (ND) as −𝑙𝑜𝑔(𝜌) and fitness differences (FD) as 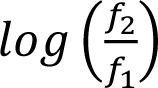. While other definitions of ND and FD are more flexible (Spaak *et al*. 2023) these definitions allow us to compare ND and FD on the same scale, and to recover the ecologically intuitive condition for species coexistence, i.e., the ND between a pair of species must be greater than their FD for them to coexist.

### Adding temperature sensitivity to model parameters

For each mechanistic process from the MacArthur model (i.e., *r_k_*, *K_k_*, *c_ik_*, *v_ik_*, *m_i_*; Eq. 4), we modeled temperature sensitivity using an exponential Arrhenius-form function:

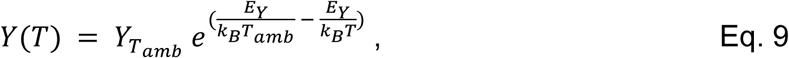

where *Y* is any parameter of the consumer-resource model (Eq. 4), *T* is the environmental temperature in Kelvin (K), *T_amb_* is the ambient temperature, here set to 10 °C (= 283.15 K), 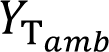 is the rate of the process at the ambient temperature, *E_Y_* is the exponential rate of increase in process rate with increasing temperature (Fig. 1; hereafter temperature sensitivity), and *k_B_* is Boltzmann’s constant (eV/K). The temperature sensitivity parameter, *E_Y_*, is equivalent to the activation energy parameter from chemistry’s Boltzmann-Arrhenius model, which has been applied to ecological rates within the Metabolic Theory of Ecology (Brown *et al*. 2004).

By standardizing environmental temperature, *T*, relative to ambient temperature, *T_amb_*, we isolate the effects of 15 °C warming, rather than the particular range of temperatures employed for our analyses (10 – 25 °C). While it is known that most individual- and population-level temperature responses are unimodal (Amarasekare & Savage 2012; Kingsolver 2009) and that exponential models typically overpredict the temperature sensitivity of processes (Knies & Kingsolver 2010; Michaletz & Garen 2024; Pawar *et al*. 2016), many studies have focused on the rate of increase in the exponential portion of the temperature-response curve (Fig. 1B, 1C) to model scaling effects of temperature (Bernhardt *et al*. 2018a; Gibert *et al*. 2022; Gilbert *et al*. 2014). As such, we used an exponential model to describe warming-induced shifts in competition when both species were in this exponentially increasing portion of their thermal niche, and where the effects of temperature were non-stressful (sub-optimal). We obtained temperature dependence estimates for each parameter from posterior distributions (Fig. 2) generated from an empirical data synthesis for each parameter (see *Supplemental Methods*).

**Figure 2.**
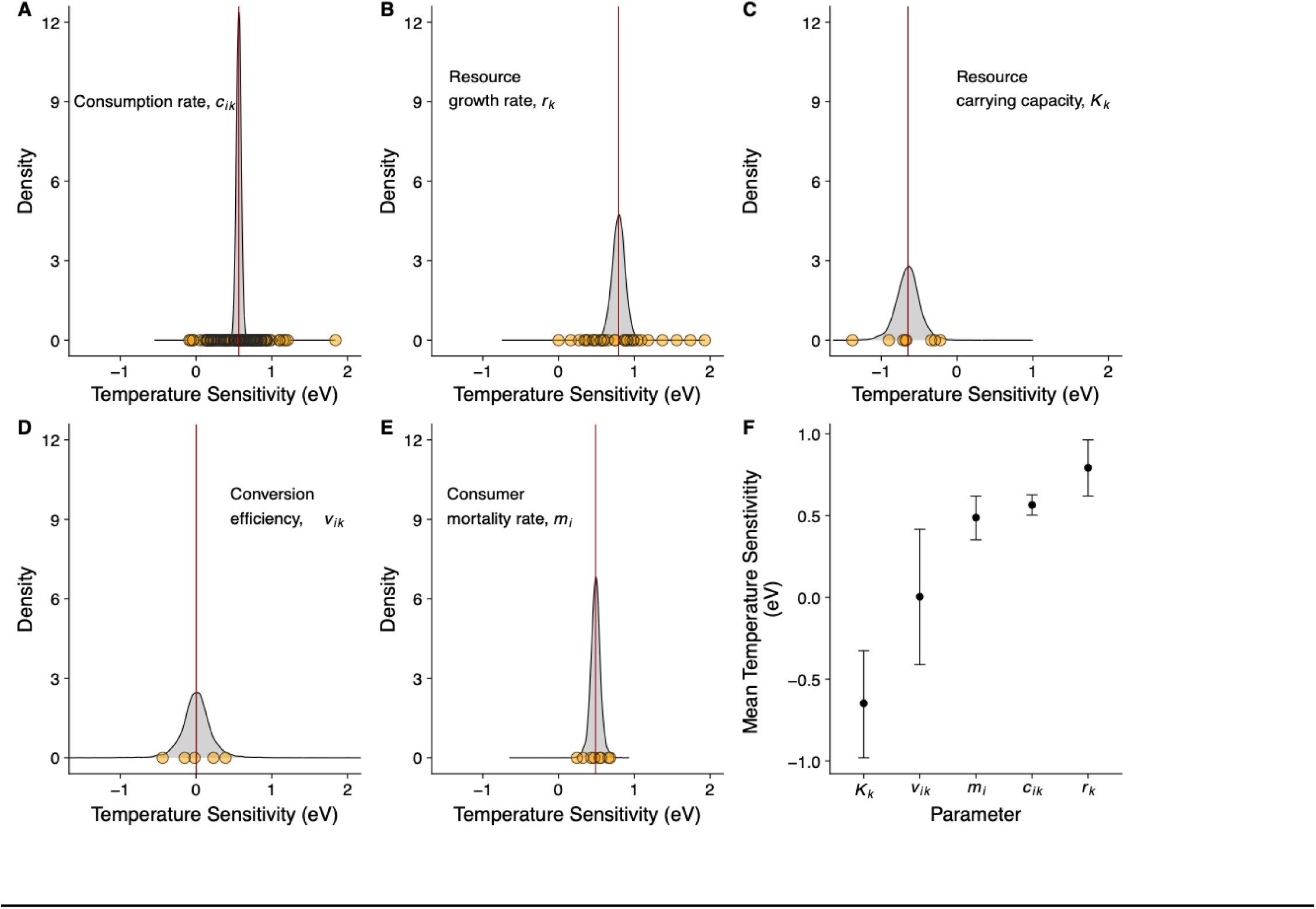
Posterior distributions for temperature sensitivity of consumer-resource model parameters. Panels A-E show published empirical observations for each parameter in the consumer resource model (Eq. 4) in orange circles and posterior distributions informed by these observations in grey density plots. Red lines indicate the mean of the posterior distribution. Panel F shows the means and 95% credible intervals (CIs) for each parameter.

### Simulating effects of warming on competition

#### Model starting conditions

We answered our three questions by simulating competition across a 15-degree temperature gradient (10°C to 25°C) between consumer species pairs that co-occur or coexist under ambient temperatures (*T_amb_* = 10°C). All simulations presented in the main text had the same starting conditions at *T_amb_*, captured by the intercepts in the Arrhenius models (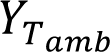 ; Eq. 9, Table S1) that comprise the temperature-dependent consumer-resource model (Eq. 4): at *T_amb_*, species were predicted to coexist and consumer species pairs have symmetrical resource preferences, so that each consumer specialized to the same degree on one of two substitutable resources (given by *c_ik_* and *v_ik_*; Table S1). At *T_amb_* the preferred resource of *N_2_* (here, *r_a_*) had a faster growth rate than the preferred resource of *N_1_* (here, *r_b_*). This set of starting conditions placed our species pairs on the boundary between coexistence and competitive exclusion of *N_1_* by *N_2_* (Fig. 3). Other starting conditions for competing consumers that co-exist under ambient temperature conditions were explored in supplementary analyses (Table S1, Figs. S2, S3, S4). Although 15°C of warming is unrealistic for changes in average temperatures from a climate change perspective, we used this large thermal gradient to demonstrate our results clearly. Applying a more realistic thermal gradient (5 °C) generated qualitatively similar results, but with less contrast (Fig. S5).

**Figure 3.**
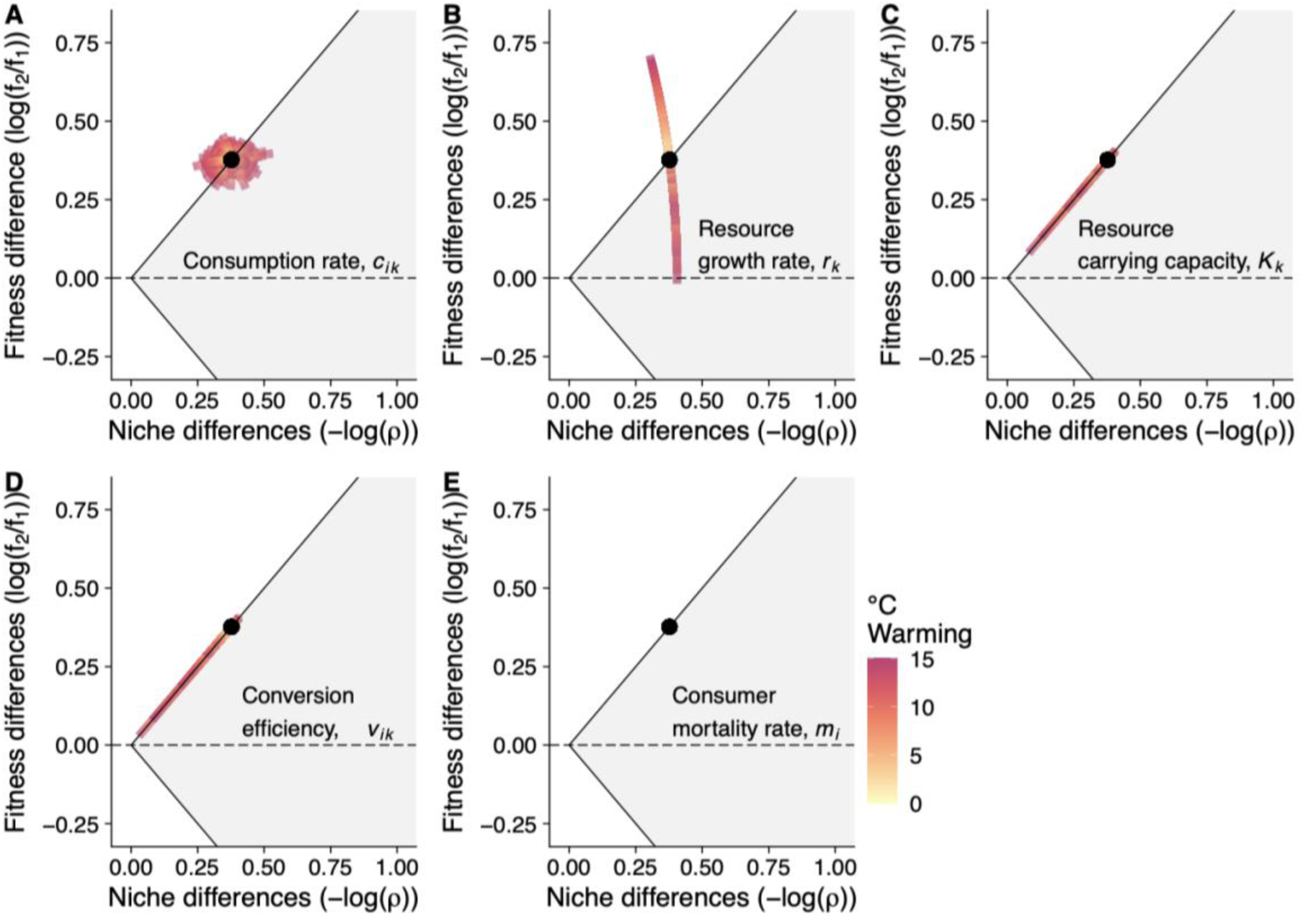
Effects of temperature sensitivity in each model parameter on competition with warming. In each panel, temperature sensitivities of the focal parameter are drawn randomly from the parameter’s empirical distribution (Fig. 2) and then the system is warmed 15°C, while temperature sensitivities for all other parameters are held at 0. This is repeated 500 times and each simulated warming path is plotted. All simulations begin at the same ambient temperature, 10°C (black point). For all parameters except for consumption rate (i.e. panels B-E), the number of temperature sensitivities drawn for each simulation is two -- one for each consumer or resource species (Fig. 1A). This results in perfectly overlain trajectories for each simulation: note different lengths of the temperature gradient path, which ends with a dark red color. For consumption rate, in panel A, there are four temperature sensitivities drawn for each simulation, corresponding to one temperature sensitivity per resource, for each species (Fig. 1A). As a result, warming trajectories do not fall perfectly on top of one another. The shaded region in the plots shows where species can coexist, i.e., where 𝑁𝐷 > |𝐹𝐷|, and the unshaded regions show where one species competitively excludes the other, i.e., 𝑁𝐷 < |𝐹𝐷|.

**Figure 4.**
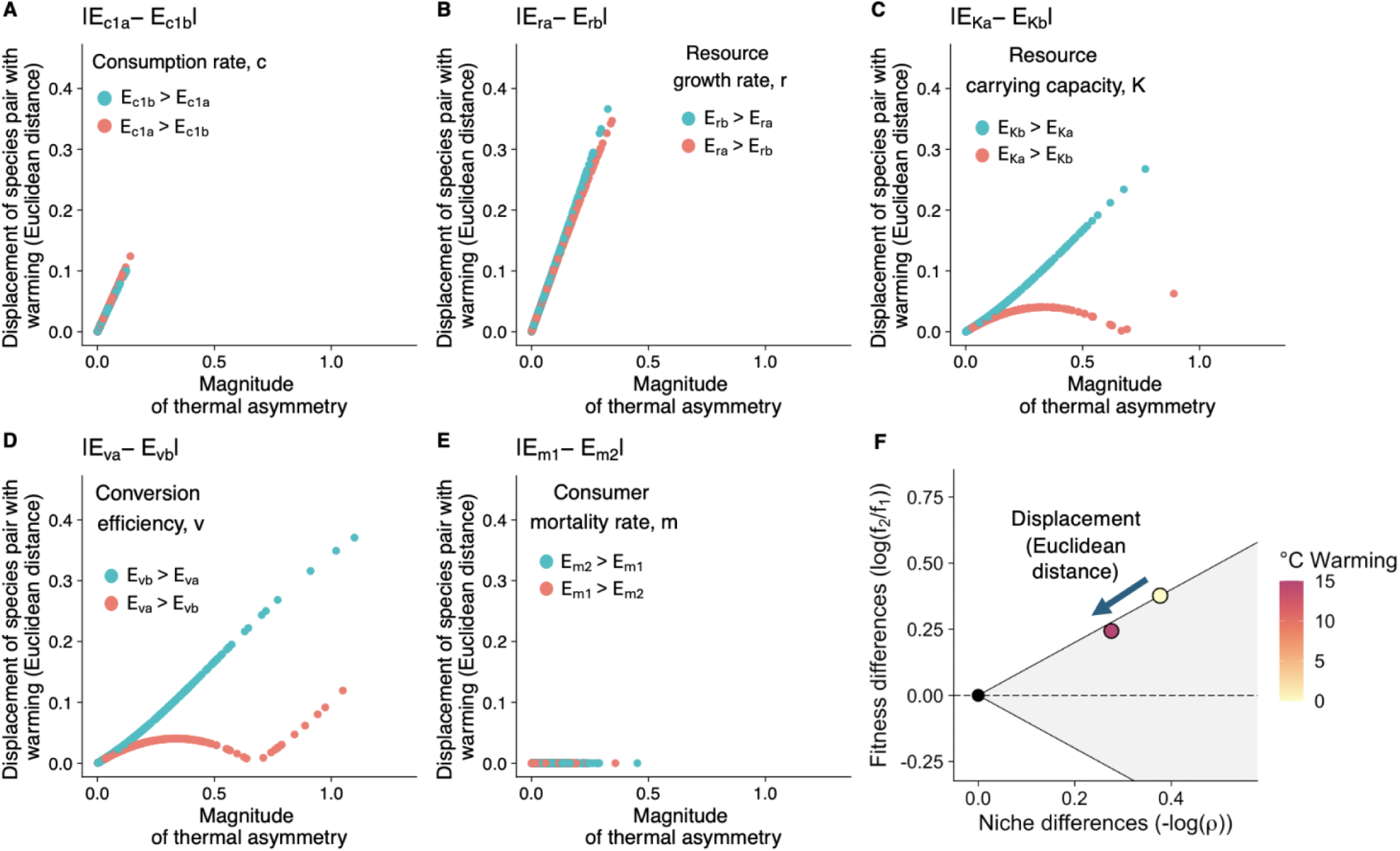
Shifts in competition with warming are related to intra-process thermal asymmetries. Panels A-E show displacement of species pairs with warming as a function of thermal asymmetries in each parameter in the consumer-resource model, and panel F depicts displacement of species pairs with warming on canonical niche difference-fitness difference axes. Thermal asymmetries are given as the absolute value of the difference between two draws of temperature sensitivities (one for each consumer or resource species) in each simulation and displacement is given as the Euclidean distance between a species pair’s starting position and its position after 15°C warming. For each panel (A-E), only the focal temperature sensitivity parameter was allowed to vary; all other temperature sensitivity parameters were fixed at 0. Data points correspond to the difference in temperature sensitivities drawn for each of 500 simulations, and the color of the points indicates the direction of the thermal asymmetry, i.e. which *E_Y_* value is larger. For consumption rate (panel A), we vary only two of the four possible temperature sensitivities, here species *N_1_*’s consumption rate of each resource, to display the effect of this thermal asymmetry most clearly.

#### Simulation: Questions 1 and 2

To examine the effects of each parameter’s temperature sensitivity on ND and FD (Question 1) and the effects of thermal asymmetries on competition (Question 2), we ran 500 warming simulations, varying one focal parameter at a time. For each simulation we randomly drew temperature sensitivity, *E_Y_*, values for the focal parameter from its empirical posterior distribution, and we set all other parameters’ temperature sensitivities equal to 0. After each temperature sensitivity for the focal parameter was drawn, we simulated 15°C warming and calculated ND and FD every 0.1°C, according to Eqs. 7 and 8. For resource growth rate, *r_k_*, resource carrying capacity, *K_k_*, and consumer mortality rate, *m_i_*, temperature sensitivities for each consumer or resource were drawn independently for each simulation. For consumption rate, *c_ik_,* we drew temperature sensitivities independently for each species’ consumption rate on each resource (See Fig. S6 for other draw configurations). Finally, for resource conversion efficiency, *v_ik_*, we assumed that the temperature sensitivity primarily reflects the nutritional composition of the resource, and therefore was the same for both consumers of a given resource (i.e. 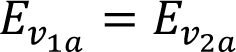).

To explore the effects of inter- and intra-process thermal asymmetries on competition (Question 2), we quantified the magnitude of the thermal asymmetries from the temperature sensitivities drawn for each simulation and examined the associated shift in ND–FD space (See *Supplemental Methods*). We quantified the magnitude of thermal asymmetry as the absolute difference between the values drawn for the focal parameter in each simulation (e.g. 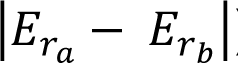). We then quantified the shift in ND–FD space as the Euclidean distance between the model’s starting position and the position after 15°C simulated warming.

#### Simulation: Question 3

To identify general effects of warming on competition (Question 3), we used the same set of starting conditions as in Questions 1 and 2 (Table S1), but imposed temperature sensitivity on all model parameters simultaneously, then simulated warming over a 15°C thermal gradient. For each simulation (N = 500), we drew temperature sensitivity values, *E_Y_*, for all parameters from their respective distributions, then calculated ND and FD at each 0.1°C step. We quantified effects of warming on competition based on the distance between the species’ starting ND and FD values (at *T_amb_*, 10°C) and their median ND and FD values at the final temperature (25 °C).

Data and code for analyses are available at DOI: 10.5281/zenodo.17253649.

## Results

### Empirically-derived temperature sensitivity of parameters

We synthesized consumer-resource parameters’ temperature sensitivity estimates from 58 studies, representing 79 ectothermic consumer or resource species. Represented species included heterotrophic and autotrophic as well as unicellular and multicellular organisms (Table S2, Fig. S7). We found considerable variation within and across temperature sensitivity estimates, *E_Y_* (Fig. 2; Table S3), suggesting both inter-process and intra-process thermal asymmetries were present in our synthesized dataset. Posterior distributions and 95% credible intervals for the temperature sensitivities of resource growth rate, *r_k_*, consumption rate, *c_ik_*, and consumer mortality rate, *m_i_*, were positive, indicating that these rates increase with temperature. Posterior means and 95% credible intervals were: 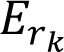 = 0.79 eV (0.62, 0.96 eV; 30 observations); 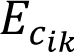 = 0.56 eV (0.50, 0.63 eV; 105 observations); 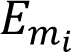 = 0.49 eV (0.35, 0.62 eV; 8 observations; Fig. 2A, 2B, 2E). Overlapping 95% credible intervals across these three parameters suggests no inter-process thermal asymmetry among them (Fig. 2F). Contrastingly, resource carrying capacity, *K_k,_* had a negative temperature sensitivity, 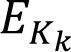 = -0.65 eV (−0.98, -0.33 eV; 8 observations; Fig. 2C), indicating that carrying capacities decrease with warming. This negative temperature sensitivity was statistically distinct from that of consumption rate, resource growth rate, and mortality rate (Fig. 2F). Finally, the posterior distribution for the temperature sensitivity of conversion efficiency had a 95% credible interval that spanned both negative and positive values and was centered near 0: 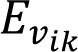 = 0.0036 eV (−0.41,0.42 eV; 5 observations; Fig. 2D). Credible intervals suggest inter-process thermal asymmetries between conversion efficiency, consumption rate and resource growth rate, but not between resource carrying capacity and conversion efficiency or mortality rate (Fig. 2F).

### Question 1: How do temperature sensitive processes underlying competition affect niche and fitness differences?

For competing consumer species that co-exist under ambient temperature conditions, warming the system caused every process except mortality to have large effects on niche and/or fitness differences (ND, FD; Fig. 3). Effects of warming on resource consumption rate, *c_ik_*, drove variation in both ND and FD (Fig. 3A), whereas effects of warming on resource growth rate, *r_k_*, primarily drove variation in FD (Fig. 3B). We found that temperature sensitivity in these two processes had the largest potential to affect competitive outcomes in our model, because their temperature sensitivity can move the species pair along a trajectory that is far from parallel (and in some cases close to perpendicular) to the coexistence boundary as the system warms. In contrast, effects of warming on resource carrying capacity, *K_k_*, and conversion efficiency, *v_ik_*, drove variation parallel to the coexistence boundary (Fig. 3C, 3D, S3). In our focal analysis, where species pairs underwent warming from a starting position on the coexistence boundary, temperature-dependent carrying capacity and conversion efficiency drove the species towards increased functional similarity (i.e. toward the neutral point).

In supplemental analysis of co-existing species pairs with different resource preferences at ambient temperature, and therefore different starting points in ND—FD space (Table S1), effects of temperature-sensitivity in resource consumption rate, *c_ik_*, resource growth rate, *r_k_*, and mortality, *m_i_*, were similar, while effects of temperature-sensitivity in resource carrying capacity, *K_k_*, and resource conversion efficiency, *v_ik_*, differed (Fig. S2C-D, S3C-D). Specifically, temperature-sensitive resource carrying capacity and conversion efficiency had qualitatively different effects on ND and FD within the 15°C temperature gradient when consumers’ resource preferences were sufficiently different at the ambient temperature. That is, when warming amplified the competitive dominance of one consumer species over the other (e.g. when *N_1_* had a stronger preference for its preferred resource than *N_2_*, and conversion efficiency of that preferred resource increased with warming), those consumers increased in ND over the 15°C temperature gradient, and the species’ competitive trajectory pointed away from the neutral point (Fig. S3D).

Across all analyses temperature-sensitive of consumer mortality rate only affected FD. Competitive dynamics were highly insensitive to variation in the temperature sensitivity of mortality, as indicated by the absence of visible shifts in Fig. 3E (see also Fig. 4E).

### Question 2: How do thermal asymmetries contribute to shifts in competition with warming?

We found that competing species that responded identically to warming (i.e. had identical temperature sensitivities for all parameters) experienced no change in competitive dynamics with warming (Fig. S8). That is, without intra-process thermal asymmetries, inter-process thermal asymmetries did not affect competitive dynamics. However, when species differed in their temperature sensitivities for a given process, competitive dynamics shifted with warming. Within each parameter, larger differences in temperature sensitivities between competing species or between their resources (intra-process thermal asymmetries) were associated with larger shifts in competition, i.e. larger Euclidean distances between the species’ “start” point at the ambient temperature and the “end” point after 15°C warming (Fig. 4, S9). For consumer mortality rate, *m_i_*, changes in position with warming were too small relative to that of other parameters to be visible in our analysis (Fig. 4E).

For each parameter except consumption rate, warming created two distinct trajectories that moved in opposite directions away from the ambient temperature point (Fig. 4B-E). Which trajectory a species pair moved along depended on the relative value of the two focal temperature sensitivities, or the directionality of the intra-process thermal asymmetry (Fig. 4B-E). For resource growth rate and consumer mortality rate, these trajectories moved monotonically toward the region of NFD space where a consumer’s preferred resource was more abundant with warming, or where consumer mortality was minimized, though the latter effect was too small to observe for most species pairs. For example, if the temperature sensitivity for growth rate was higher for resource *a* than for resource *b*, then, when the system was warmed, the coexistence trajectory moved monotonically toward the region of competitive dominance for the species that preferentially consumes resource *a*, here *N_2_* (e.g. Fig. 3B, 4B). Interestingly, the two trajectories for carrying capacity and conversion efficiency comprised one monotonic trajectory and one non-monotonic trajectory, both of which pointed ultimately toward neutrality (Fig. 3C-D, 4C-D, S3). However, large differences in ambient-temperature resource preferences between competing species drove qualitative changes to these trajectories over the 15 °C warming gradient (Table S1, Fig. S3).

### Question 3: How does warming affect competition via effects on niche and fitness differences?

For consumer species existing at the boundary of coexistence and competitive exclusion, the temperature sensitivity of all processes underlying competition generally drove decreases in both ND and FD with warming (Fig. 5A, yellow dot to magenta dot). The majority of species pairs decreased in ND (58%) or FD (58%) and the median decreases in ND and FD were similar in magnitude (5.5% decrease and 9.7% decrease, respectively; Fig. 5C-D). Declines in ND and FD with warming were associated with decreasing resource carrying capacity and resulting declines in all competition coefficients (Fig. S10). This led to a general trend of competitive dynamics shifting toward neutrality with warming (Fig. 5A). The distribution of shifts in competition with warming was skewed, with very large shifts being rare (Fig. 5B). The magnitude of shift in competition was positively related to the magnitude of intra-process thermal asymmetries (Table S4). This result is reinforced by analysis of competition between species at the extremes of the observed temperature sensitivity distributions (top and bottom 5%), which showed much larger shifts in competition with 15°C warming (Fig S11).

**Figure 5.**
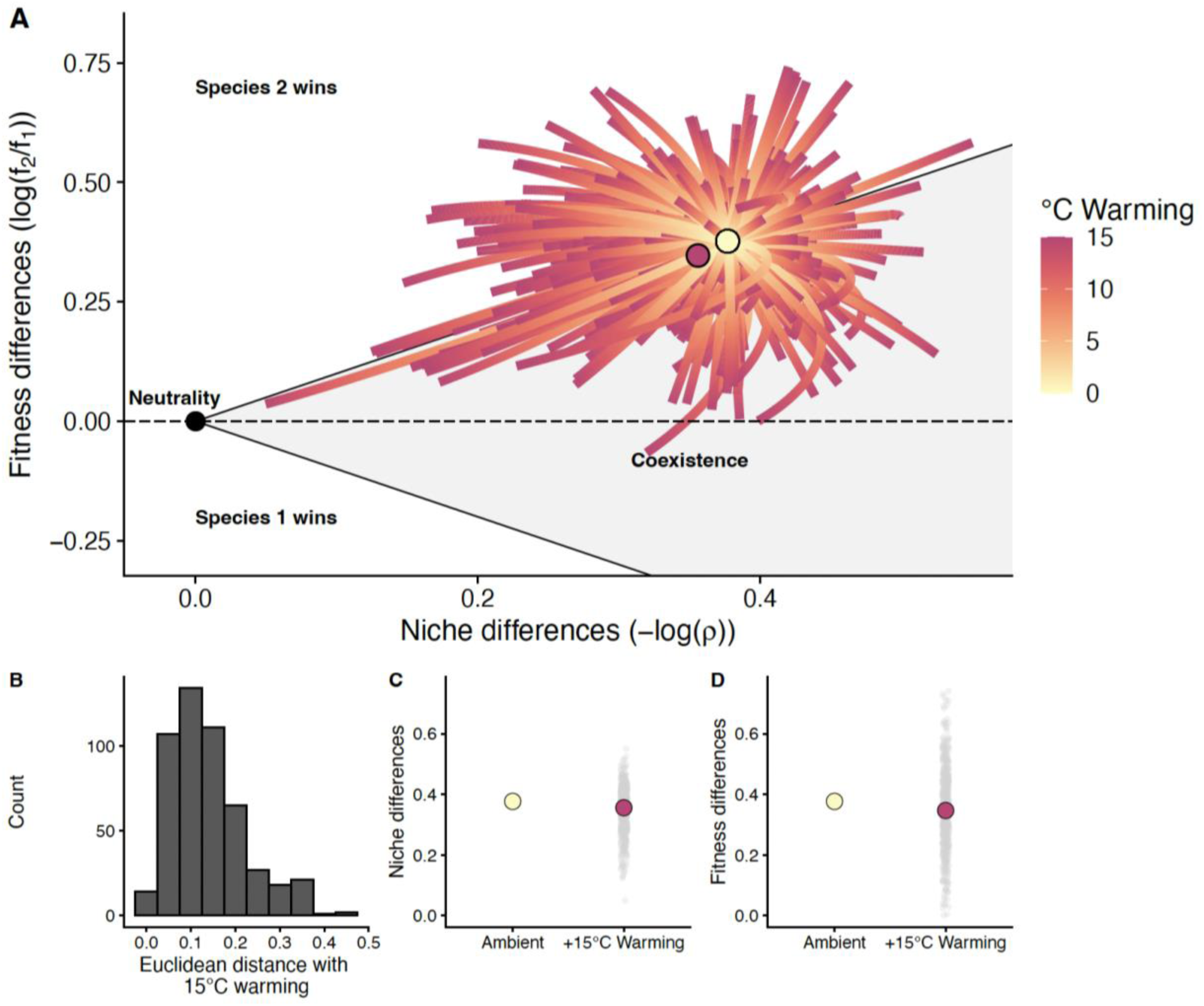
Effects of warming on competition. All parameters are drawn from their distributions and then warmed 15°C, and this simulation is repeated 500 times. A) Simulation path for each draw of parameter values from the ambient temperature, 10°C (yellow dot), to 25°C (15°C warming). The median position along both axes after 15°C warming is plotted as a magenta dot. B) Distribution of Euclidean distances between the position at the ambient temperature and after 15°C warming across 500 simulations. C) Shifts in niche differences with warming. The median niche difference values (magenta dot) and the niche difference values for each simulation (grey dots) after 15°C warming are shown compared to the value at the ambient temperature (yellow dot). D) Shifts in fitness differences with warming, displayed as in panel C.

While some competitive dynamics became less neutral (i.e. increased in ND and FD) under 15°C warming, simulating extreme warming revealed that trajectories that initially moved away from neutrality bent back toward neutrality with sufficient warming (Fig. S10, S12). These results were robust to different consumer and resource traits and numbers of resource species, but the amount of warming required to shift the median effect toward neutrality under these different conditions varied (Figs. S1-S4).

## Discussion

In this study, we drew from two prominent theories in ecology to address a central question in global change biology: how does warming affect competition for shared resources? Using empirically-derived temperature sensitivities of the processes underlying competition, we found three general patterns in how simulated competition responded to warming. First, we found that warming tended to reduce both fitness differences and niche differences. This resulted in a general trend of warming causing species pairs to move toward neutrality. Next, we found that warming-induced increases in mortality had surprisingly little effect on these shifts in competition with warming. Finally, we found that species pairs that underwent the largest shifts in competition with warming had highly asymmetric temperature sensitivities in the processes underlying competition. Together, these findings suggest that understanding how communities will change with warming requires an understanding of how both population vital rates and resource use traits change with temperature, and how variable these traits are within a community.

While much of the work seeking to understand biodiversity effects of climate change has focused on temperature-induced changes in fitness differences (e.g. growth rates; Ehrlén & Morris 2015; Kordas *et al*. 2011; Merow *et al*. 2014), here we show that understanding the effects of warming on competition requires an equal consideration of niche differences. As expected, resource growth rate had a large effect on competition in our analysis. This effect was near orthogonal to the coexistence boundary, occurring primarily via shifts in fitness differences and causing resource growth rate to affect competitive hierarchies (by moving species from one region of ND—FD space to another) under warming. However, warming is unlikely to affect resource growth rate in isolation, and when warming was allowed to affect the full suite of processes underlying competition, species pairs generally also decreased in niche differences with warming (Fig. 5). These shifts were driven primarily by the effects of warming on consumption rate, resource carrying capacity, and consumer conversion efficiency – all of which determine patterns of relative resource use among species. Of these processes, warming effects on consumption rates also had the capacity to alter competitive hierarchies (Fig. 3A). Together, these results add to a limited body of literature that suggests that temperature effects on niche differences play a key role in how warming affects communities (Adler *et al*. 2012).

Warming-induced changes to mortality rates have been a focal point of climate modelling and thermal physiology (Deutsch *et al*. 2008; Sunday *et al*. 2014), and have been shown to have effects on competition and, specifically, on fitness differences (Song *et al*. 2019). In our analysis, despite exponentially increasing consumer mortality rates across the thermal gradient (Amarasekare & Savage 2012; Simon & Amarasekare 2024) effects of warming on mortality rate had almost no effect on competition. This is due to the independence of consumer mortality rate from effects of interspecific competition in consumer-resource models (Eq. 6; Song *et al*. 2019), and is consistent with other work suggesting that mortality rates alone cannot drive shifts in coexistence (Chesson 2000; Smith & Amarasekare 2018). This suggests that, when species are living at temperatures far below their thermal optimum, and biological processes underlying competition are increasing exponentially with warming, quantifying effects of warming on mortality may be less important than previously thought.

Within Modern Coexistence Theory, competition is considered neutral when there is an absence of both niche differences and fitness differences; biologically this means that species are functionally equivalent (Adler *et al*. 2007). In our analysis, warming had a general tendency to shift competing species towards neutrality, regardless of species thermal traits under ambient temperatures (Fig. 5, S1-S5). In our simulations, this tendency to move toward neutrality as temperatures increased was related to the exponential decrease in resource carrying capacity with warming (Fig. 2, S10). As consumers became increasingly limited by reduced resource supply, intra- and inter-specific competition weakened, and consumers thus became more ecologically similar to one another. This result is consistent with theoretical (Hubbell 2001; Lande 1993) and empirical (Schreiber *et al*. 2022; Siepielski *et al*. 2010) work showing that smaller population sizes are associated with increased neutrality and stochasticity. Additionally, our finding that systems trend toward neutrality regardless of competitors’ traits under ambient conditions is consistent with recent work suggesting that that there may be characteristic trajectories for how competition changes under extrinsic influences, rather than bespoke shifts based on interspecific trait differences (Pastore *et al*. 2021).

While our finding that warming generally makes competition more neutral is independent of the model’s starting conditions, we found that the specific resource-use and thermal traits of species influence the amount of warming required to shift competition toward neutrality (Fig. 5, Figs. S1-S4). That is, for some consumer pairs, an increase in neutrality may not be realized within a biologically plausible range of temperatures because the amount of warming required to create this effect might be large. In addition, if warming exceed species’ thermal optima for one or more of the processes underlying competition, then competitive trajectories may diverge from those predicted here (Fig. S13). This is relevant in systems where species’ thermal optima are close to mean environmental temperatures, and may also be relevant in systems with highly variable environmental temperatures, which at times exceed thermal optima (Amarasekare & Johnson 2017; Casas Goncalves & Amarasekare 2021; Smith & Amarasekare 2018).

Under changing thermal regimes, which types of species pairs are likely to experience large competitive shifts remains an open question. While most species simulated in our study experienced relatively small shifts in competitive dynamics with warming, species pairs that experienced large shifts had large intra-process thermal asymmetries (Fig. 4, Table S4, Fig. S9). While a considerably body of work has described how inter-process thermal asymmetries can drive cross-scale responses to warming (e.g., Dell *et al*. 2014; Gibert *et al*. 2022; Gilbert *et al*. 2014; Lopez-Urrutia *et al*. 2006), in our analysis, inter-process thermal asymmetries required accompanying intra-process thermal asymmetries in order to affect competition. For example, a higher temperature sensitivity of resource growth rate than conversion efficiency cannot, on its own, cause a shift in competition between species (Fig. S8): there must also be interspecific variation in the temperature sensitivities of the growth rates or conversion efficiencies of the different resources to affect competition. More generally, we found that consumer species (or their resources) that have the most divergent thermal traits will undergo the largest shifts in competition with warming. Previous studies have demonstrated the effects of warming on competition, usually based on differences in thermal optima among species to generate effects of warming on competition (Simon & Amarasekare 2024; Smith & Amarasekare 2018). Here, we show for the first time that general effects of warming on competition can emerge from interspecific differences in suboptimal thermal traits alone, demonstrating how warming changes competition even in conditions that are considered thermally benign for all species considered.

## Conclusions

Making general predictions for the effects of warming on communities has been a major challenge in ecology. The Metabolic Theory of Ecology, with its focus on the temperature sensitivity of metabolic processes across biological scales, and Modern Coexistence Theory, through its conceptual framework for species competition, have both advanced our understanding of ecosystem responses to environmental change. However, differences in the domains of these two fields, one of which focuses on interspecific differences and the other which focuses on macroecological differences in key metabolic processes as drivers of change, have prevented their integration to date. Here we integrated concepts from both fields and combined this with empirical thermal trait data to develop new, general theory for how competition changes with warming and advance a more coherent understanding of global change. The next step will be to test these predictions empirically. Doing so will facilitate better linkages between general patterns in biological responses to warming and context-specific stewardship strategies under rapid environmental change.

## Authorship Statement

KED, JRB, TNG, PJK, PLT, and MIO conceptualized the study. JRB collected the data. KED, JRB and PJK, developed software and analyzed the data. All authors contributed to writing, editing and revising the manuscript.

## Data Accessibility Statement

Data and code for analyses are available at <u>DOI: 10.5281/zenodo.17253649</u>.

## Acknowledgements

JRB, TNG, and MIO were supported by Discovery Grants from the Natural Science and Engineering Research Council of Canada, and KD was supported by a Centre for Ecosystem Management Postdoctoral Fellowship. We thank Megan Szojka for helpful feedback on the manuscript.

## Notes

### Competing Interest Statement

The authors have declared no competing interest.

### Summary of Updates

We contextualized our work with other work on temperature-dependent competition, added several new analyses demonstrating the robustness of our results (primarily the trend of decreasing niche and fitness differences with warming), and clarified a major mechanism behind these results.

