## Supplemental Information for "General predictions for the effects of warming on competition"

Supplemental Methods

**Data Synthesis: Estimating temperature sensitivity of model parameters**

To employ the best estimates of temperature sensitivity, *E_Y_*, for each mechanistic parameter in our model, we compiled data from the literature on the temperature sensitivity (Figure 1B and 1C) of consumption rate, *c_ik_*, conversion efficiency, *v_ik_*, resource carrying capacity, *K_k_*, mortality, *m_i_,* and growth rate, *r_k_*, for ectothermic consumer species and ectothermic resource species. Our objective was not to perform a new exhaustive systematic review of the literature, but to synthesize evidence for temperature sensitivities from a broad, representative and taxonomically unbiased portion of the literature. We achieved this goal by drawing on an existing systematic review (e.g., Dell *et al.* 2011) and a new synthesis of relevant datasets. Our dataset contains 157 observations of temperature sensitivities from 79 species of ectotherms from freshwater, marine and terrestrial habitats. We assembled this synthetic dataset by searching the primary literature using Google Scholar for empirical datasets describing the temperature sensitivity of biological processes and rates listed above using combinations of search terms that describe the biological processes of interest (i.e. rates of consumption, mortality, growth, as well as conversion efficiency and carrying capacity) and terms describing the temperature effects (i.e. temperature, warming, activation energy). We conducted our searches using combinations of search terms for biological processes (“consumption rate”, “conversion efficiency”, “carrying capacity”, “growth rate”, “population growth rate”, “intrinsic rate of increase”, “mortality rate”, “lifespan”) and temperature effects (“temperature”, “activation energy”, “temperature dependence”) and combinations of these terms (e.g. “consumption rate AND activation energy”, or “mortality rate AND temperature”). We used a broad range of search terms to obtain as taxonomically unbiased a sample of the literature as possible. Because temperature sensitivities can be reported in many different ways in the literature, we used a backward and forward snowballing method (also known as citation chaining) to identify additional relevant papers. We included data from existing published compilations of activation energies such as (2011). Some studies returned in our search reported lifespan as a function of temperature, and in those cases, we calculated mortality rate as 1/lifespan, following McCoy and Gillooly (2008). For each estimate of temperature sensitivity, we report the maximum and minimum temperatures over which the temperature sensitivity was calculated. We included multiple types of measurements for our biological processes. For example, for the growth rate of the resource, we included estimates of intrinsic rate of increase and specific growth rate. We restricted our dataset to studies that included at least four experimental temperatures to allow for accurate estimation of temperature sensitivities (Pawar *et al.* 2016) and to experiments conducted in controlled laboratory conditions. We extracted metadata and values for biological processes and temperatures from text, tables, or from figures using WebPlotDigitizer (Rohatgi 2022). We included studies that reported activation energies estimates and three studies that provided raw data sufficient for us to calculate activation energies using a linearized exponential model, which we did using ordinary least squares regression in R. Our initial search yielded 251 observations of temperature sensitivities for our focal biological processes; we retained 157 observations after filtering our dataset according to the inclusion criteria listed above. To account for unevenness in sampling and uncertainty around temperature sensitivity estimates in the represented studies, we used the empirical estimates of temperature sensitivity to generate an empirical distribution of temperature sensitivities for each parameter. We did so by fitting an intercept-only linear regression to empirical estimates of *E_Y_* for each parameter, and then Gibbs sampling the posterior distribution (burn-in = 1000 iterations; N = 10000 samples). We ran this analysis using the ‘MCMCregress’ function in the ‘MCMCpack’ (Version 1.7.1; Martin *et al.* 2011) package in R.

For all parameters, we pooled all observations and drew from across the full posterior distribution in our simulations. We then assumed that resource dynamics are much faster than consumer dynamics and used a timescale separation technique to simplify the consumer-resource model. While combining data from diverse sources could violate the fast-resources assumption by pairing fast consumer data with slow resource data, this does not affect equilibrium outcomes (Case & Casten 1979), so should not affect our results. Our dataset combines observations from aquatic and terrestrial environments, heterotrophic and autotrophic resources, as well as multicellular and unicellular resources. We did not have sufficient coverage within groups to enable reliable parameter estimation for each temperature sensitivity parameter within groups, so we pooled data across groups in our simulations. Because we pooled data across systems to generate posterior distributions, is possible that simulations included combined data across groups, for example temperature sensitivities from aquatic consumers with terrestrial resources. However, mitigating this issue would require sufficient empirical observations within each of these groups across all parameters to construct distributions for each group-parameter combination, which we were not able to do with our dataset due to limitations on within-group sample sizes

**Interpreting effects of warming on competition**

After calculating niche and fitness differences at each temperature step, we displayed the results from this analysis on orthogonal fitness and niche difference axes, so that each species pair is represented as a single point coordinate in ND—FD space. Over the temperature gradient, the 150 points per species pair form a thermal trajectory travelling away from the model’s start point.

Following the notation of modern coexistence theory, the criterion for species coexistence in a two-species Lotka–Volterra model can be written as $\rho<\frac{f_{2}}{f_{1}}< \frac{1}{\rho}$ ; after log-transformation and using our definition for niche and fitness differences, the criterion is equivalent to $ND > \left| FD \right|$, which matches the intuition that coexistence requires species niche differences to be greater than their fitness differences. The parameter space of niche–fitness differences is then divided into a shaded region, where species can coexist, and unshaded regions where one species competitively excludes the other (Figure 3). Whether warming-induced movement through ND–FD space (here, ‘warming trajectory’) crosses from one region into another is dependent on the starting conditions of the system, so we instead focus on the length of the trajectory and directional trends in niche and fitness differences with warming.

**Derivation of the generalized equilibrium consumer-resource model (Equation 4):**

$\frac{dN_{i}}{dt}=\sum_{k\in\left\{ a,b \right\}} v_{ik}c_{ik}R_{k}N_{i}-m_{i}N_{i}$	Main text eq. 1

$\frac{dR_{k}}{dt}=r_{k}R_{k}\left( 1-\frac{R_{k}}{K_{k}} \right)-\sum_{i=1}^{2} c_{ik}R_{k}N_{i}$ ,	Main text eq. 2

Assuming fast dynamics and solving for quasi-equilibrium, $\hat{R_{k}}$ gives:

$\hat{R_{k}}=\frac{K_{k}}{r_{k}}\left( r_{k}-\sum_{i=1}^{2} c_{ik}N_{i} \right)$ ,	Main text eq. 3

which is a function of consumer abundances. Substituting this expression back into the equation of consumer dynamics, we arrive at the following for the two-resource system, where:

$$\frac{dN_{i}}{dt}=N_{i}\left\{ v_{ia}c_{ia}\left[ \frac{K_{a}}{r_{a}}\left( r_{a}-c_{ia}N_{i}-c_{ja}N_{j} \right) \right]+v_{ib}c_{ib}\left[ \frac{K_{b}}{r_{b}}\left( r_{b}-c_{ib}N_{i}-c_{jb}N_{j} \right) \right]-m_{i} \right\}$$

$=N_{i}\left[ \left( v_{ia}c_{ia}K_{a}+v_{ib}c_{ib}K_{b}-m_{i} \right)-\left( v_{ia}\frac{K_{a}}{r_{a}}c_{ia}^{2}+v_{ib}\frac{K_{b}}{r_{b}}c_{ib}^{2} \right)N_{i}-\left( v_{ia}\frac{K_{a}}{r_{a}}c_{ia}c_{ja}+v_{ib}\frac{K_{b}}{r_{b}}c_{ib}c_{jb} \right)N_{j} \right]$,

This can be generalized to a multiple resource system as:

$\frac{dN_{i}}{dt}N_{i}=\left[ \left( \sum_{k\in\{a,b\}} v_{ik}c_{ik}K_{k}-m_{i} \right) - \left( \sum_{k\in\{a,b\}} v_{ik}\frac{K_{k}}{r_{k}}c_{ik}^{2} \right)N_{i}-\left( \sum_{k\in\{a,b\}} v_{ik}\frac{K_{k}}{r_{k}}c_{ik_{jk}^{c}} \right)N_{j} \right]$

Main text eq. 4

Supplemental Tables

**Table S1. Starting conditions for each competition scenario.** Each scenario is implemented in the model (Eq. 4) through a set of values at the ambient temperature, i.e., $Y_{T_{amb}}$ values in the temperature-sensitivity function (Eq. 9), e.g c_1a_ = $c_{{1a}_{T_{amb}}}$.

| **Scenario description**  **(relevant figures)** | **Consumer-resource preferences** | | | **Other parameter values** |
| --- | --- | --- | --- | --- |
| 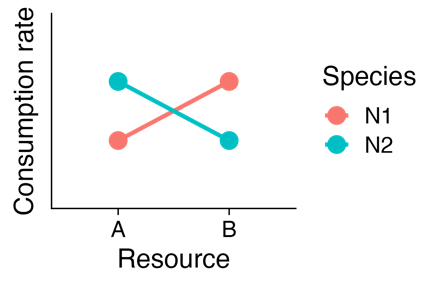1. Symmetrical resource preference + uneven resource growth rates at *T_amb_* (Figures 3, 4, 5, S5, S6, S8-S13) |  | ***R_a_*** | ***R_b_*** | r_a_ = 1, r_b_ = 0.5  K_a_ = 2000, K_b_ = 2000  m_1_ = 0.01, m_2_ = 0.01 |
|  | ***N_1_*** | c_1a_ = 0.5  v_1a_ = 0.5 | c_1b_ = 1  v_1b_ = 1 |  |
|  | ***N_2_*** | c_2a_ = 1  v_2a_ = 1 | c_2b_ = 0.5  v_2b_ = 0.5 |  |
| 2. Symmetrical resource preference + even resource growth rates at *T_amb_* (Figure S2)  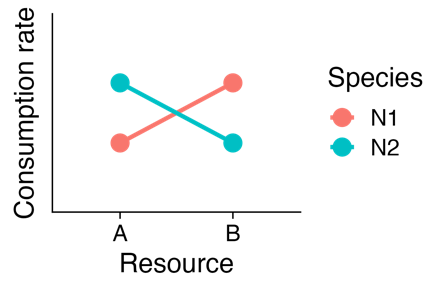 |  | ***R_a_*** | ***R_b_*** | r_a_ = 0.5, r_b_ = 0.5  K_a_ = 2000, K_b_ = 2000  m_1_ = 0.01, m_2_ = 0.01 |
|  | ***N_1_*** | c_1a_ = 0.5  v_1a_ = 0.5 | c_1b_ = 1  v_1b_ = 1 |  |
|  | ***N_2_*** | c_2a_ = 1  v_2a_ = 1 | c_2b_ = 0.5  v_2b_ = 0.5 |  |
| 3. Asymmetrical resource preference + even resource growth rates at *T_amb_* (Figure S3)  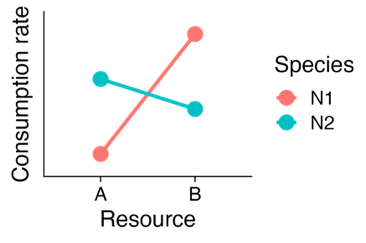 |  | ***R_a_*** | ***R_b_*** | r_a_ = 0.5, r_b_ = 0.5  K_a_ = 2000, K_b_ = 2000  m_1_ = 0.01, m_2_ = 0.01 |
|  | ***N_1_*** | c_1b_ = 0.1  v_1b_ = 0.1 | c_1b_ = 0.9  v_1b_ = 0.9 |  |
|  | ***N_2_*** | c_1a_ = 0.6  v_1a_ = 0.6 | c_1b_ = 0.4  v_1b_ = 0.4 |  |

**Table S2. Demographics of the dataset in terms of cellularity and trophy**

| **Parameter** | **Cellularity** | **N** | **Trophy** | **N** |
| --- | --- | --- | --- | --- |
| Consumption rate | Unicellular | 1 | Autotrophic | 0 |
|  | Multicellular | 104 | Heterotrophic | 105 |
| Conversion efficiency | Unicellular | 2 | Autotrophic | 0 |
|  | Multicellular | 3 | Heterotrophic | 5 |
| Mortality rate | Unicellular | 0 | Autotrophic | 0 |
|  | Multicellular | 8 | Heterotrophic | 8 |
| Resource carrying capacity | Unicellular | 7 | Autotrophic | 2 |
|  | Multicellular | 1 | Heterotrophic | 6 |
| Resource growth rate | Unicellular | 17 | Autotrophic | 13 |
|  | Multicellular | 13 | Heterotrophic | 17 |

**Table S3. Observed ranges compared to 95% credible intervals for the temperature dependence of each parameter.**

| **Parameter** | **95% Credible Interval** | **Observed range (n)** |
| --- | --- | --- |
| *E_cik_* | 0.50, 0.63 | 0.09, 1.8 (105) |
| *E_rk_* | 0.62, 0.96 | 0, 1.9 (30) |
| *E_mi_* | 0.35, 0.62 | 0.24, 0.68 (8) |
| *E_Kk_* | -0.98, -0.33 | -1.38, -0.22 (8) |
| *E_vik_* | -0.41, 0.42 | -0.44, 0.39 (5) |

**Table S4. Partial regression results for relationship between thermal asymmetries in consumer or resource parameters and displacement of species pairs in the main text simulation (Figure 5).** Displacement is measured as the Euclidean distance between the species position in NFD space at 10 °C (reference temperature) and at 25 °C. Significant relationships (p<0.05) are given in bold.

| **Term** | **Estimate** | **Std Error** | **t-value** | **p-value** |
| --- | --- | --- | --- | --- |
| Intercept | 0.020 | 0.008 | 2.36 | 0.019 |
| **\|E_rb_ – E_ra_\|** | **0.773** | **0.032** | **24.1** | **<0.001** |
| E_c_ik_ |  |  |  |  |
| \|E_c_1b_ – E_c_2a_\| | -0.095 | 0.100 | -0.947 | 0.344 |
| \|E_c_1b_ – E_c_1a_\| | 0.157 | 0.097 | 1.62 | 0.106 |
| \|E_c_1b_ – E_c_2b_\| | -0.011 | 0.107 | -0.103 | 0.918 |
| **\|E_c_2b_ – E_c_2a_\|** | **0.268** | **0.096** | **2.79** | **0.005** |
| \|E_c_2b_ – E_c_1a_\| | -0.163 | 0.098 | -1.66 | 0.097 |
| \|E_c_1a_ – E_c_2a_\| | 0.032 | 0.101 | 0.315 | 0.753 |
| **\|E_Ka_ – E_Kb_\|** | **0.063** | **0.017** | **3.71** | **<0.001** |
| **\|E_va_ – E_vb_\|** | **0.127** | **0.012** | **10.3** | **<0.001** |
| \|E_m1_ – E_m2_\| | 0.008 | 0.043 | 0.185 | 0.853 |

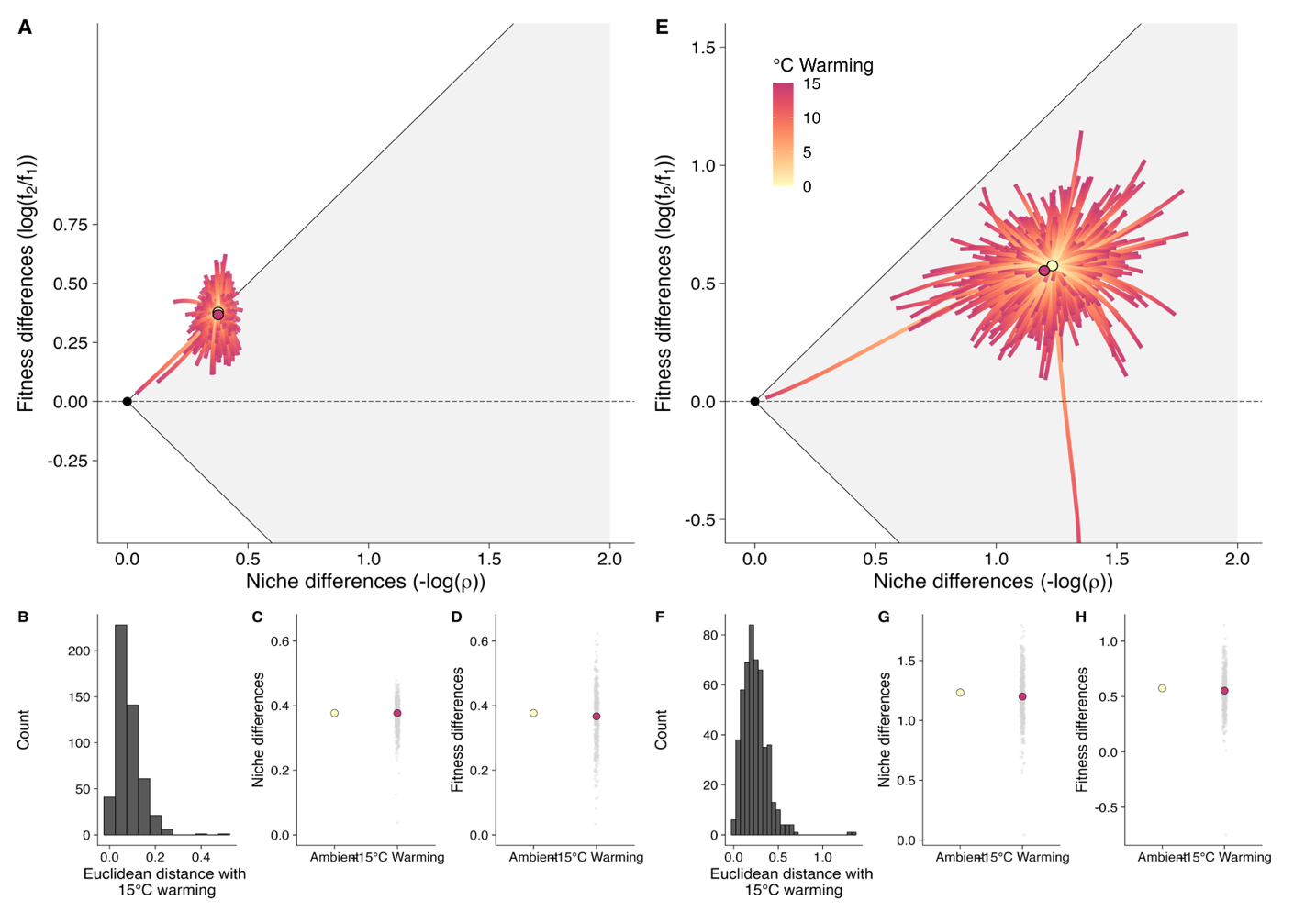
Supplemental Figures

**Figure S1. Effects of warming on competition in 2-consumer, 8-resource systems.** Panels A-D describe systems where competition is between consumer species pairs with reciprocal, symmetric resource preferences, as in the main text. Panels E-H describe systems where competition is between species pairs with asymmetric, randomly assigned resource preferences. In the symmetric scenario (Panels A-D), species 1 prefers resources a, b, c, and d, consuming each of these at a rate of 1 at the reference temperature, and consuming non-preferred resources e, f, g, and h are a rate of 0.5 at the reference temperature. Species 2 prefers resources e, f, g, and h, and consumes those at a rate of 1 at the reference temperature, while it consumes resources a-d at a rate of 0.5 at the reference temperature. Conversion efficiency is exactly equal to consumption rate at the reference temperature in all cases, and temperature sensitivities for all parameters are drawn randomly from the empirical distributions. In the asymmetric scenario (Panels E-H), consumption rates are arbitrarily assigned to each species for each resource, and conversion efficiencies match consumption rates, which assumes that consumption rates are the result of some physiological optimization for each species. In both scenarios resources a-d grow twice as fast as resources e-h at the reference temperature. Panels A and E depict the equilibrium trajectories for each species pair over a 15 °C gradient, Panels B and F depict frequency distributions for the Euclidean distance traveled by each species pair after 15 °C warming, and panels C,D and G,H depict shifts in niche differences and fitness differences for symmetric and asymmetric simulations, respectively. In panels C, D, G and H, the grey dots represent the niche and differences in each simulation at 15°C of warming, while the yellow and magenta dots are the median values of niche or fitness difference from the 500 simulations. The shift in median niche and fitness differences associated with warming can be seen by comparing the yellow and magenta dots in each panel.

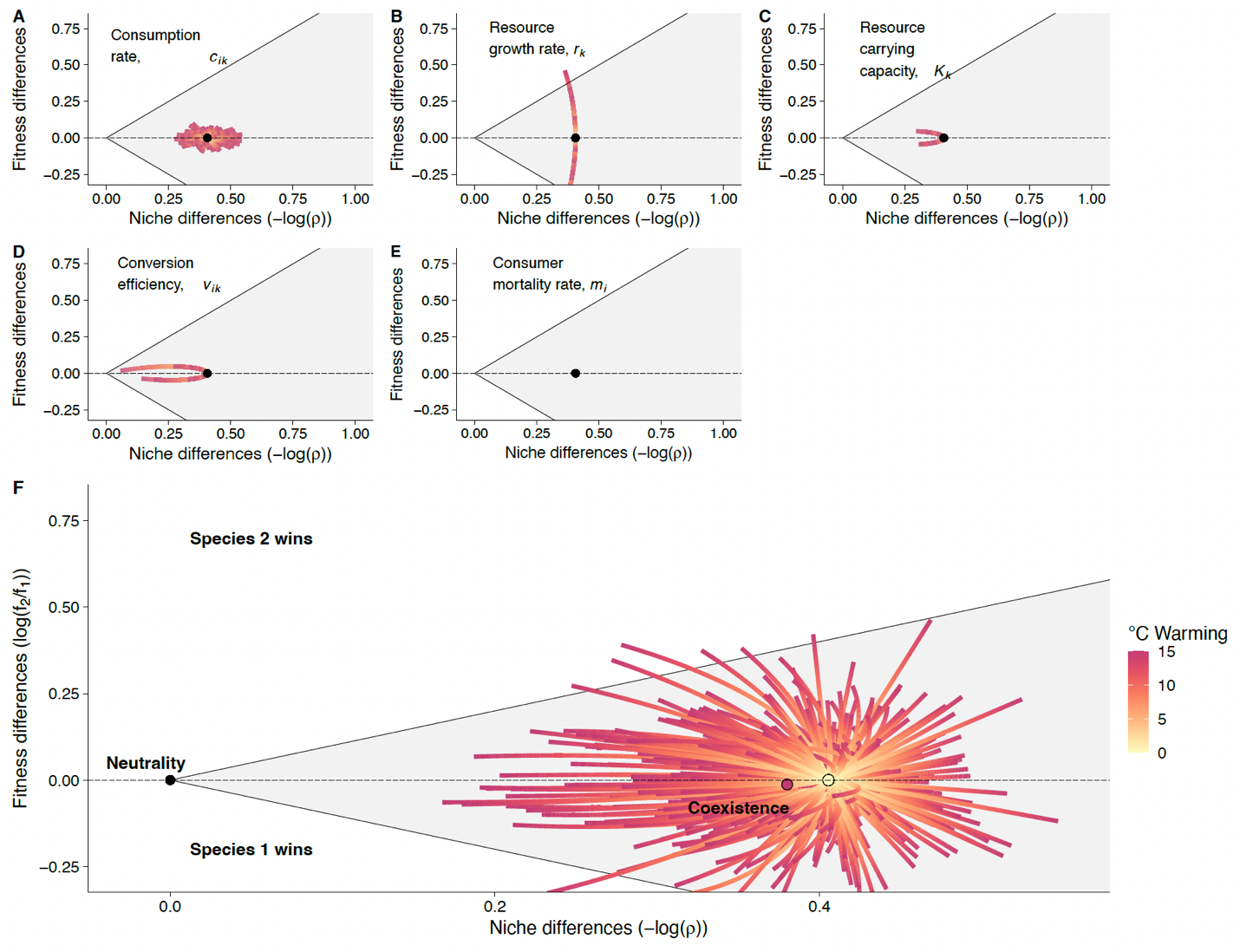

**Figure S2. Simulations of warming in species pairs with even resource preferences, where each resource has the same growth rate at ambient temperature (Table S1, Scenario 2).** The top panels show effects of warming in this system on each parameter independently (each other parameter does not vary with temperature) and the bottom panel shows the emergent effects on competition. The dominant axis of the trajectories for each parameter is similar to those in the simulation from the main text, which differs from this only in that the ambient-temperature growth rates of the two resources are unequal in that analysis. As in the main text, the yellow dot shows the start point of all species pairs (N = 500 simulations) and the magenta dot shows the median position of all species pairs after 15 °C warming.

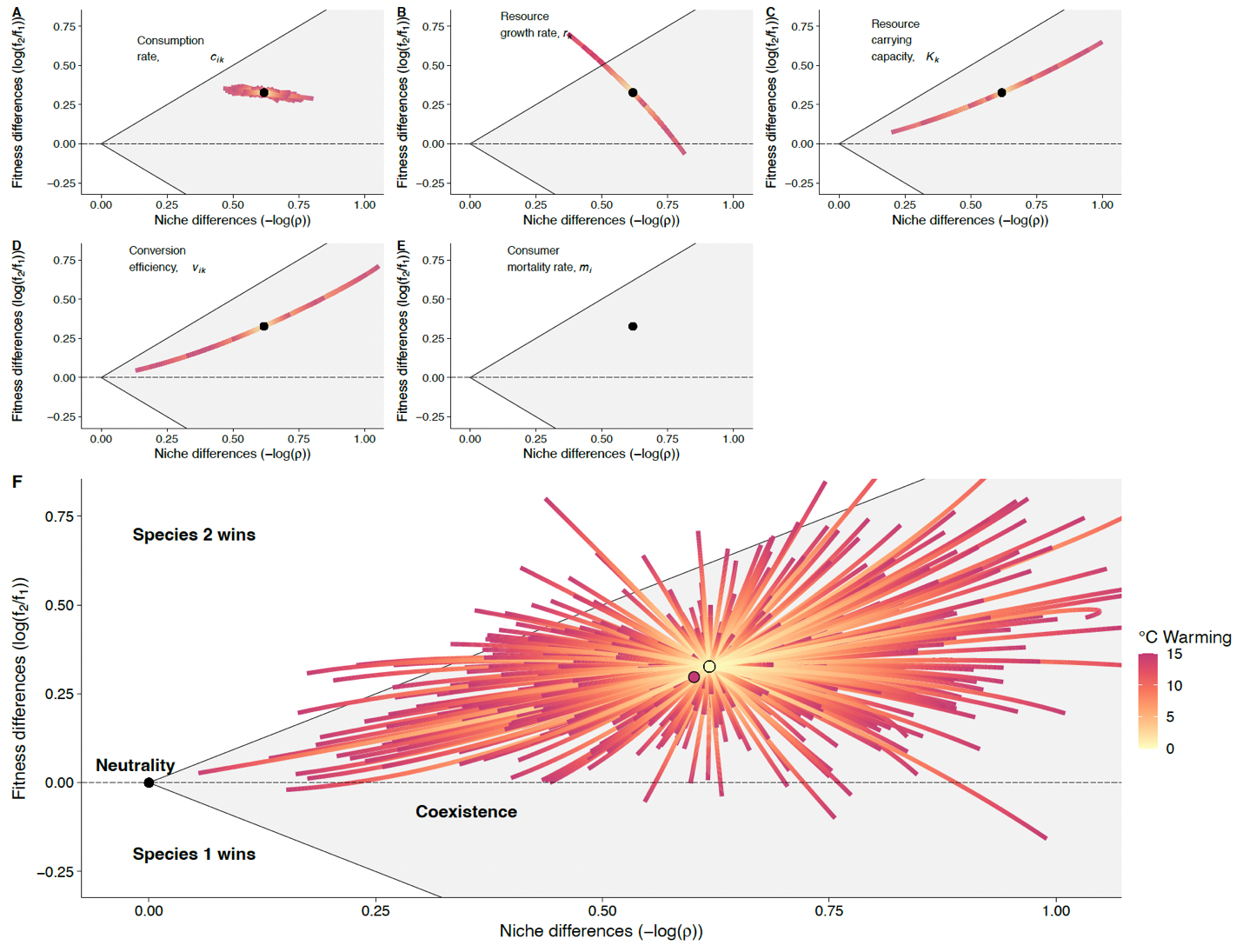
**Figure S3. Simulations of warming in species pairs with highly uneven resource preferences, where each resource has the same growth rate at ambient temperature (Table S1, Scenario 3).** The top panels show effects of warming in this system on each parameter independently (each other parameter does not vary with temperature) and the bottom panel shows the emergent effects on competition. The dominant axis of the trajectories for *c_ik_*, *r_k_*, and *m_i_* are similar to those in the simulation from the main text, while the trajectories for *K_k_* and *v_ik_* differ. When the resource preferred by the species with the weaker resource preference (Table S1) is favored by warming, either by a slower decrease in carrying capacity or higher conversion efficiency, niche differences decrease and competition shifts toward neutrality. Alternatively, when the resource preferred by the species with a stronger resource preference is favored by warming, niche differences increase and competition shifts away from neutrality. As in the main text, the yellow dot shows the start point of all species pairs (N = 500 simulations) and the magenta dot shows the median position of all species pairs after 15 °C warming.

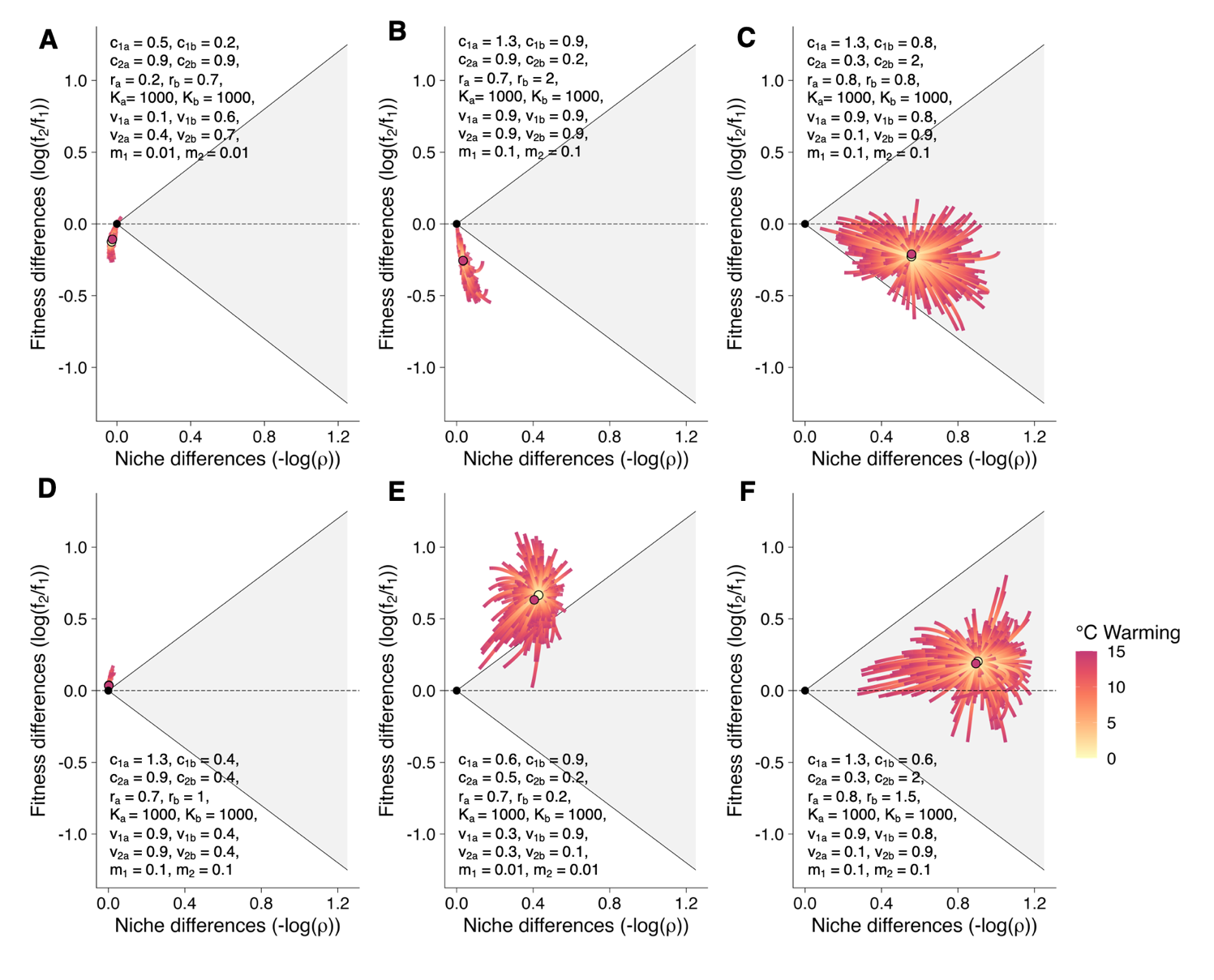

**Figure S4. Effects of warming on competition for species pairs with asymmetric resource competition.** Species are assigned arbitrary starting parameter values across *c*, *r*, and *v* at the reference temperature. The parameter values at the reference temperature are given in the insets for each panel. Temperature sensitivity values for each parameter are drawn from the empirical distributions as in the main text. In the majority of cases, across all 6 sets of starting parameter values, species’ competitive trajectories move either toward neutrality directly, or move away from neutrality but exhibit curvature back toward neutrality as they are warmed. Further, in most cases the median shift in competition with 15 °C warming (magenta point) is shifted toward the neutral (black point at 0,0) relative to the position at the reference temperature (yellow point).

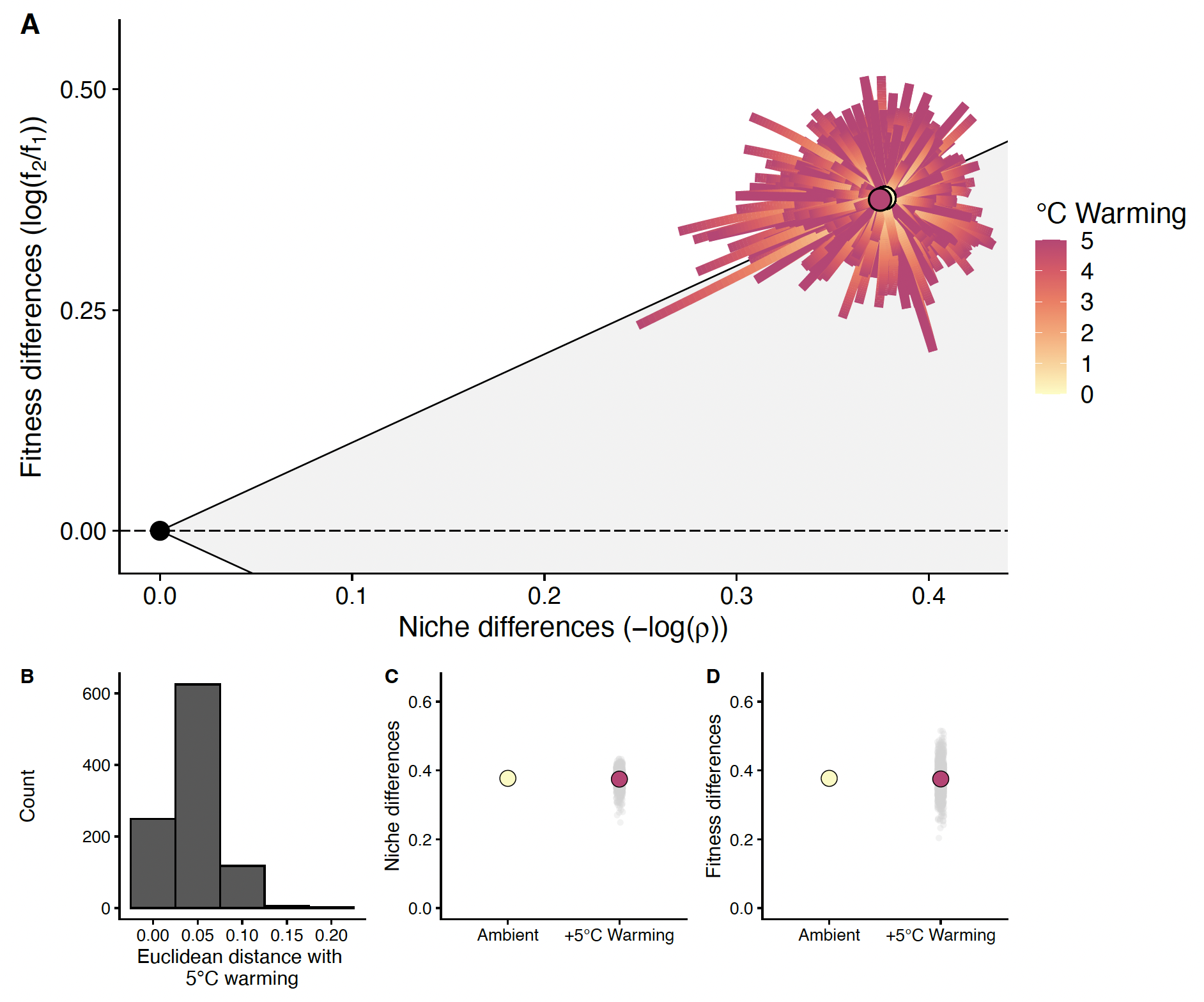

**Figure S5. Effects of 5 °C warming on competition.** All parameters are drawn from their distributions and then warmed 5 °C, and this simulation is repeated 500 times. A) Simulation path for each draw of parameter values from the ambient temperature, 10°C (yellow dot), to 15 °C (5 °C warming). The median position along both axes after 5 °C warming is plotted as a magenta dot. B) Distribution of Euclidean distances between the position at the ambient temperature and after 5 °C warming across 500 simulations. C) Shifts in niche differences with warming. The median niche difference values (magenta dot) and the niche difference values for each simulation (grey dots) after 5 °C warming are shown compared to the value at the ambient temperature (yellow dot). D) Shifts in fitness differences with warming, displayed as in panel C.

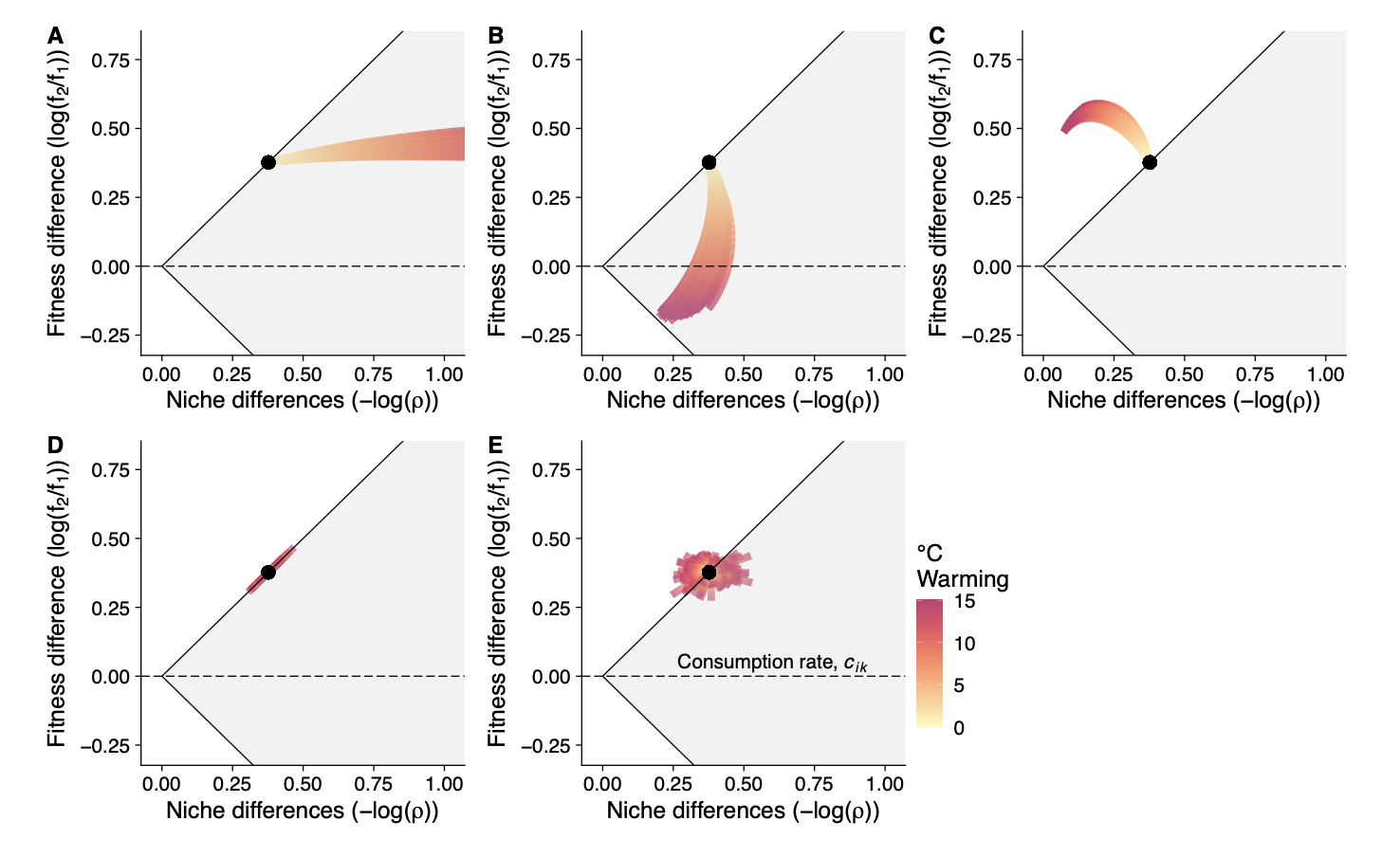

**Figure S6. Different random draw structures for the temperature sensitivity of consumption rate when all non-focal processes are not temperature sensitive.** For all panels, all processes besides consumption rate are set to temperature sensitivity = 0. A) Temperature sensitivities of consumption rates for preferred resources of each species are drawn from the empirical distribution (see Figure 2), while temperature sensitivities of consumption rates for non-preferred resources are set to 0. B) Temperature sensitivities of each species’ consumption rate of resource A are drawn from their empirical distribution, while temperature sensitivities of each species’ consumption rates of resource B are set to 0. C) Temperature sensitivities of each species’ consumption rate of resource B are drawn from their empirical distribution, while temperature sensitivities of each species’ consumption rates of resource A are set to 0. D) Temperature sensitivities of species 1’s consumption rate of each resource are drawn from their empirical distribution, while temperature sensitivities of species 2’s consumption rates of both resources are set to 0. E) All four consumption rate temperature sensitivities are drawn from their empirical distribution.

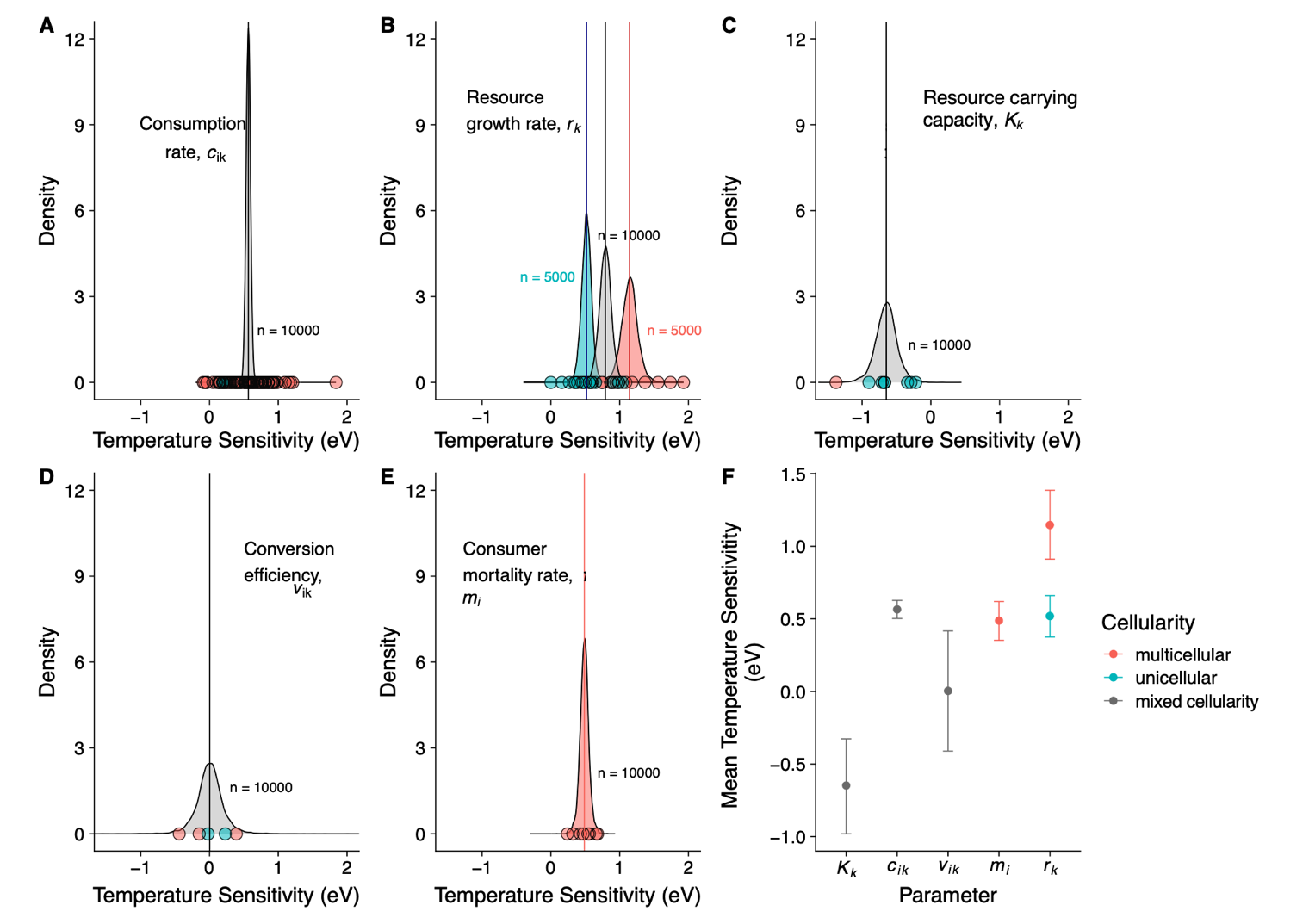

**Figure S7. Systematic variation in temperature sensitivities with cellularity.** Panels A-E show posterior distributions for either multicellular and unicellular temperature sensitivity estimates (B); or a distribution that combines multicellular and unicellular systems, where >2 observations per group were not available (A, C, D, E). The y axis indicates the posterior probability density for each temperature sensitivity value, such that high densities are related to narrow distributions and high precision, rather than an indicator of frequency (as in a histogram). The number of datapoints from the posterior distribution used to generate the density plot are indicated on each plot. Where a distribution represents one category of data, we have colored the distribution according to the cellularity it represents (B, E). Where a distribution represents data from both cellularity types, the distribution is colored grey (A, C, D). Data points along the bottom of the distribution show observations used to inform the posterior distribution, colored by the cellularity represented. Panel F shows differences in temperature sensitivity within and across parameters.

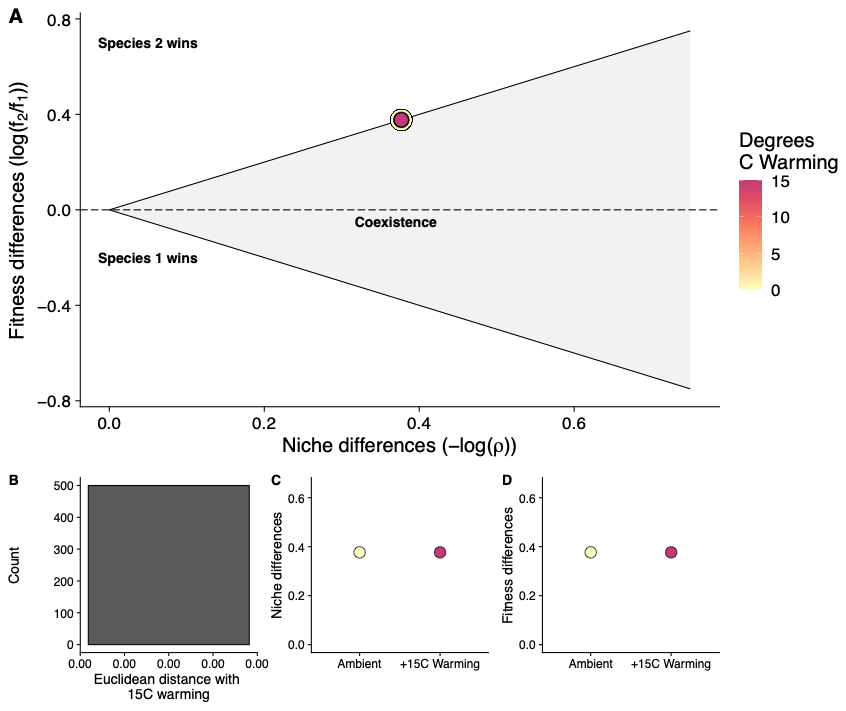

**Figure S8. Effects of warming on competition when inter-process thermal asymmetries are present, but intra-process thermal asymmetries are not.** Simulation is run identically to the main analysis (See Fig. 5 caption), but all temperature sensitivities for each process are identical between species for each draw.

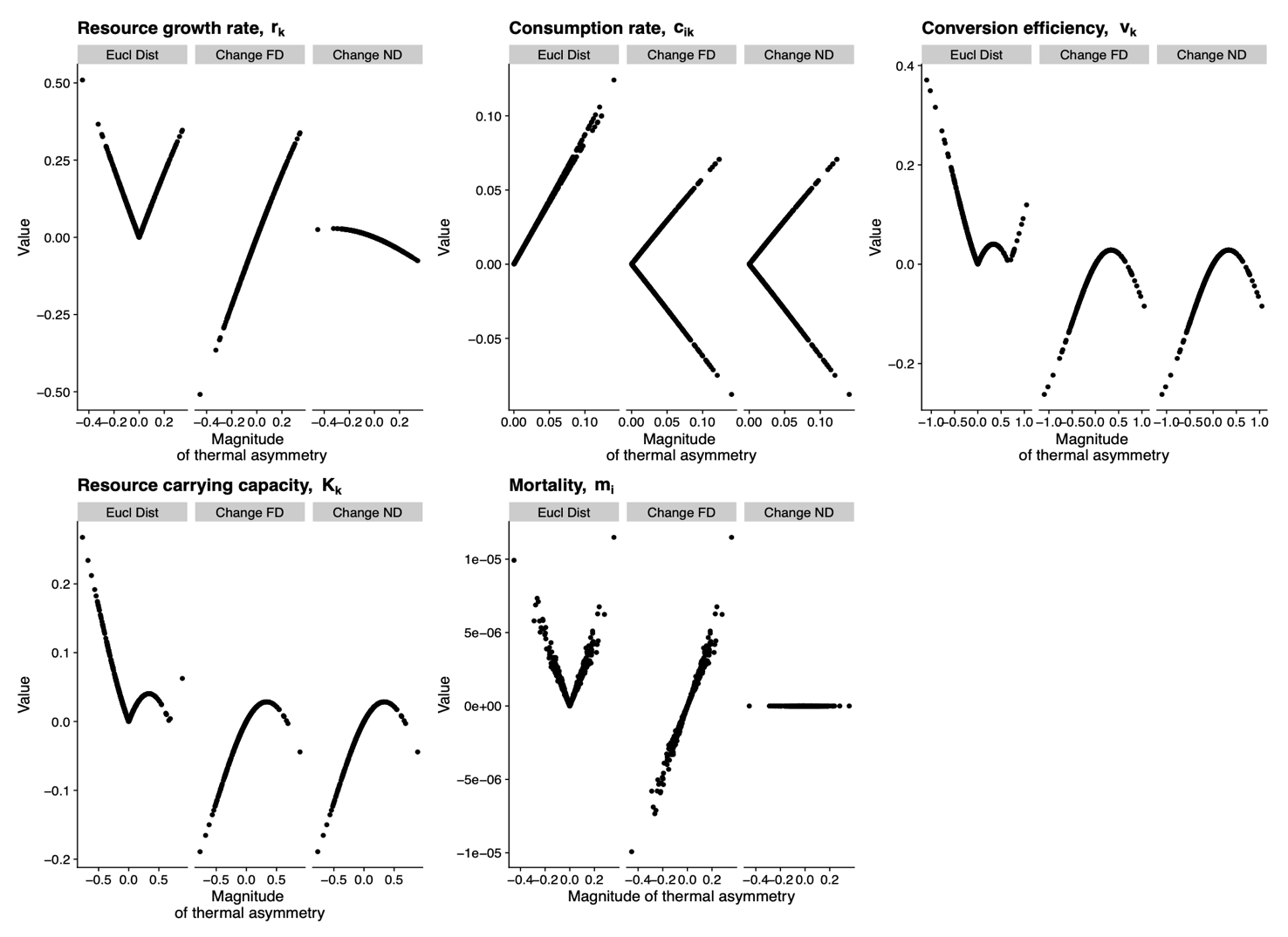

**Figure S9. Effects of thermal asymmetries in each parameter (raw values) on Euclidean distance, fitness differences (FD), and niche differences (ND).** Data are from parameter-by-parameter simulations (N = 500 per parameter), where each parameter was made temperature dependent, while all other temperature sensitivities were set to 0. Each data point is a pair of temperature sensitivity draws for the focal parameter from a single simulation of warming, and the magnitude of the thermal asymmetry is given as the difference between these two draws.

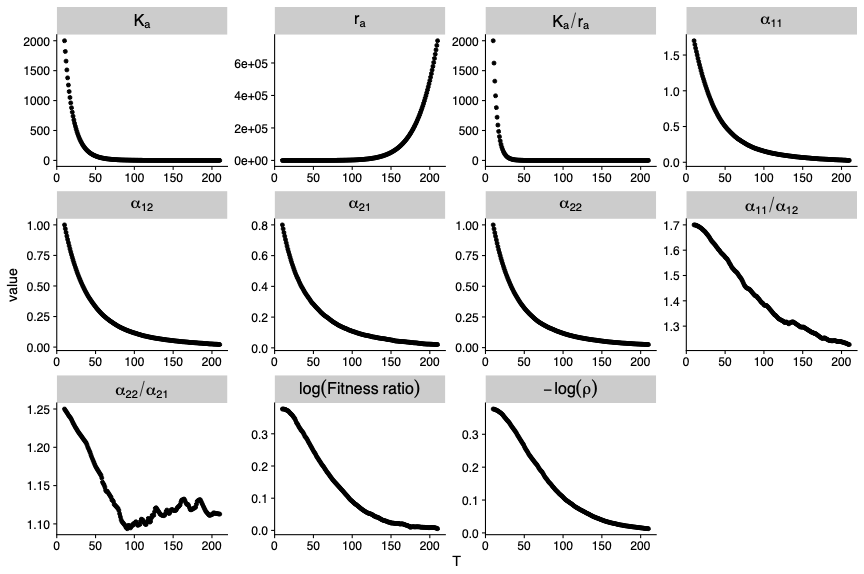

**Figure S10. Median trajectories of key model terms with 200 °C warming.** Summary statistics are calculated across 500 simulated species every 0.1 °C. Competing consumers have perfectly symmetrical resource use, as in the main text analysis (Figures 3-5). The panels labeled “log(Fitness ratio)” and “-log(rho)” correspond to the fitness differences and niche differences axes from our main analysis, respectively.

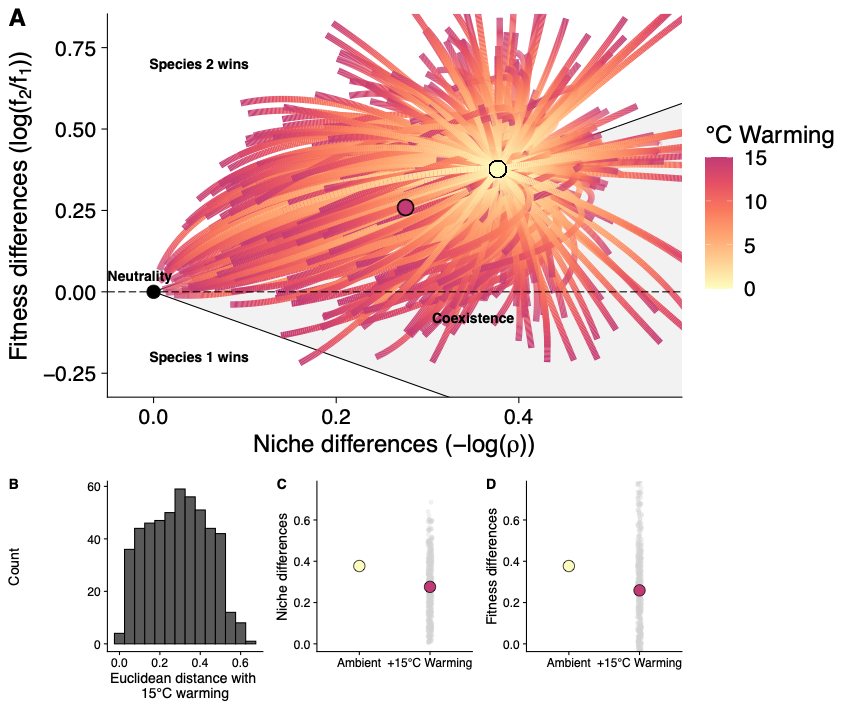

**Figure S11. Effects of extreme thermal asymmetries on competition under warming.** All parameter distributions were restricted to the top and bottom 5% of the full posterior distribution, and then temperature sensitivities for each parameter were drawn from this constrained distribution. This was repeated to generate 500 sets of species thermal traits at the extremes of the observed thermal trait distributions, then all systems were warmed 15 °C. A) Simulation path for each draw of parameter values from the ambient temperature, 10 °C (yellow dot), to 25 °C (15 °C warming). The median position along both axes after 50 °C warming is plotted as a magenta dot. B) Distribution of Euclidean distances between the position at the ambient temperature and after 15 °C warming across 500 simulations. C) Shifts in niche differences with warming. The median niche difference values (magenta dot) and the niche difference values for each simulation (grey dots) after 15 °C warming are shown compared to the value at the ambient temperature (yellow dot). D) Shifts in fitness differences with warming, displayed as in panel C.

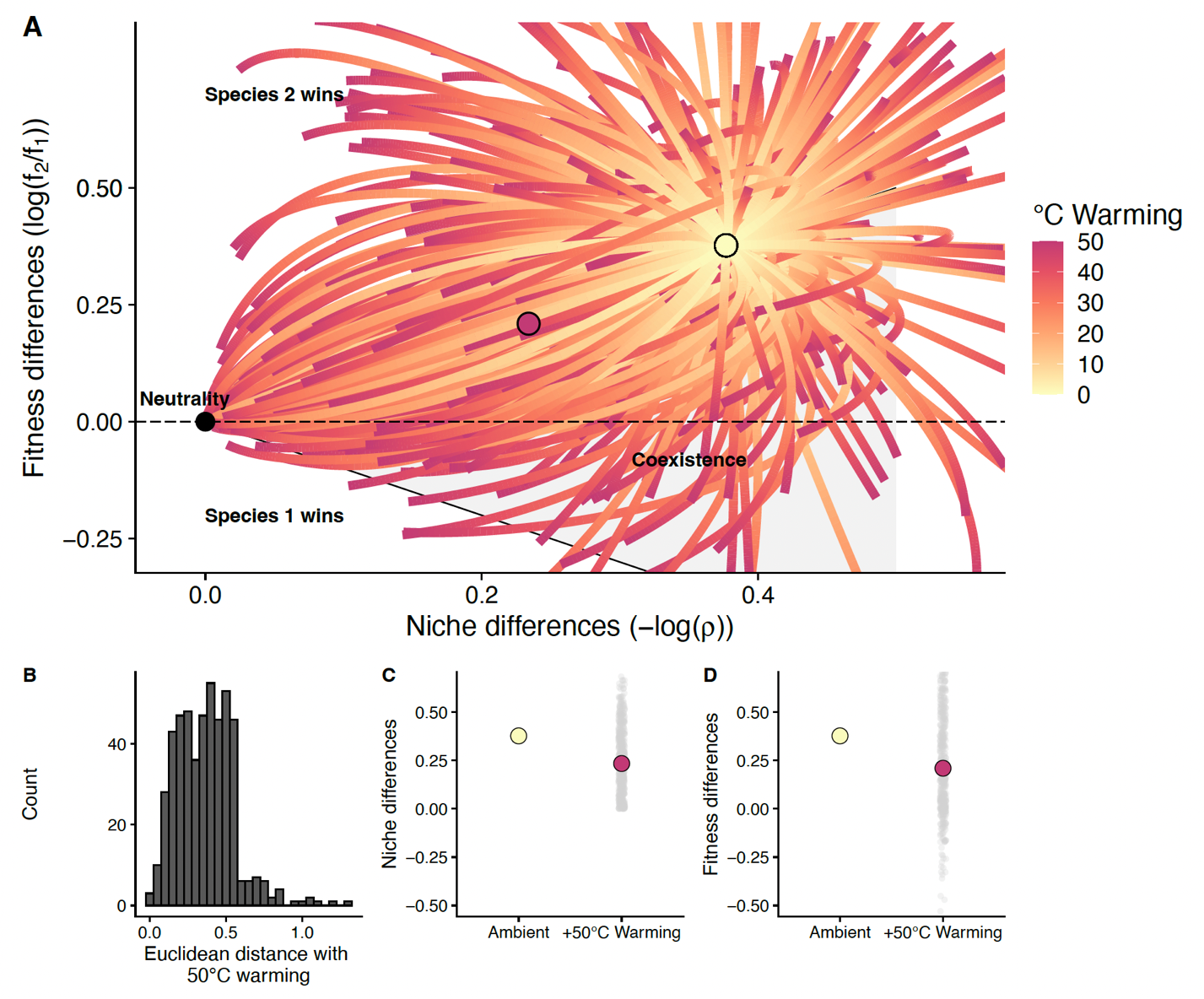

**Figure S12. Effects of extreme warming (50 °C) on competition.** All parameters are drawn from their distributions and then warmed 15 °C, and this simulation is repeated 500 times. A) Simulation path for each draw of parameter values from the ambient temperature, 10 °C (yellow dot), to 60 °C (50 °C warming). The median position along both axes after 50 °C warming is plotted as a magenta dot. B) Distribution of Euclidean distances between the position at the ambient temperature and after 50 °C warming across 500 simulations. C) Shifts in niche differences with warming. The median niche difference values (magenta dot) and the niche difference values for each simulation (grey dots) after 50 °C warming are shown compared to the value at the ambient temperature (yellow dot). D) Shifts in fitness differences with warming, displayed as in panel C.

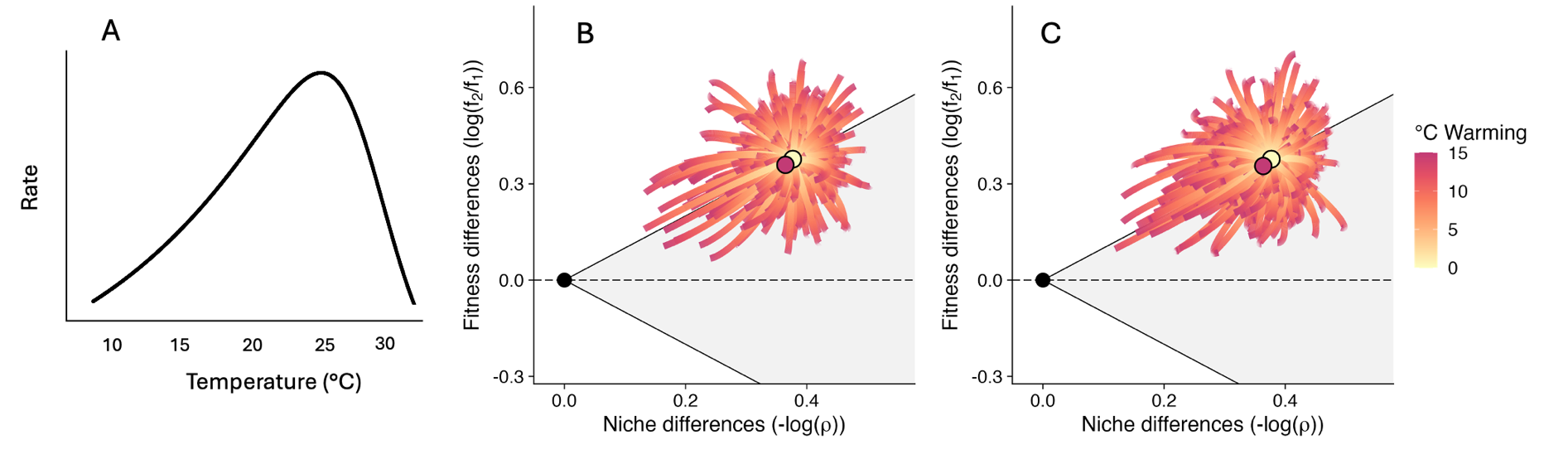

**Figure S13. Effects of unimodal temperature dependence on study results.** A) A generic unimodal thermal performance curve, which is used to characterize parameter’s temperature dependence in this analysis. B) Effects of warming on niche and fitness differences when all parameters have a unimodal temperature response that maximizes performance at 25 °C (thermal optimum, T_opt_). C) Effects of warming on niche and fitness differences when all resource parameters have a thermal optimum at 25 °C, all consumer 1 parameters have a thermal optimum at 24 °C, and all consumer 2 parameters have a thermal optimum at 23 °C.
